# Strain dropouts reveal interactions that govern the metabolic output of the gut microbiome

**DOI:** 10.1101/2022.07.25.501461

**Authors:** Min Wang, Lucas J. Osborn, Sunit Jain, Xiandong Meng, Allison Weakley, Jia Yan, William J. Massey, Venkateshwari Varadharajan, Anthony Horak, Rakhee Banerjee, Daniela S. Allende, Ricky E. Chan, Adeline M. Hajjar, Alejandra Dimas, Aishan Zhao, Kazuki Nagashima, Alice G. Cheng, Steven Higginbottom, J. Mark Brown, Michael A. Fischbach

## Abstract

The gut microbiome is complex, raising questions about the role of individual strains in the community. Here, we address this question by focusing on a functional unit within the community, the metabolic niche that controls bile acid 7α-dehydroxylation. By constructing variants of a complex defined community in which we drop out strains that occupy this niche, we explore how interactions within and between niches shape community-level metabolism. Omitting both members of the niche, *Clostridium scindens* (*Cs*) and *Clostridium hylemonae* (*Ch*), eliminates secondary bile acid production and reshapes the community in a highly specific manner: eight strains go up or down in relative abundance by >100-fold, while the remaining strains are largely unaffected. In single-strain dropout communities (i.e., a strain swap within the niche), *Cs* and *Ch* reach the same relative abundance and dehydroxylate bile acids to a similar extent. However, the effect on strains in other niches differs markedly: *Clostridium sporogenes* increases >1000-fold in the Δ*Cs* but not Δ*Ch* dropout, reshaping the pool of microbiome-derived phenylalanine metabolites. Thus, strains that are functionally redundant within a niche can have widely varying impacts outside the niche, and a strain swap can ripple through the community in an unpredictable manner, resulting in a large impact on an unrelated community-level phenotype. Mice colonized by the Δ*Cs*Δ*Ch* community show decreased liver steatosis relative to those colonized by the Δ*Ch* community, demonstrating that a single strain from the microbiome can have a substantive impact on host physiology. Our work opens the door to the mechanistic studies of the role of an individual strain on community ecology and host physiology.

## INTRODUCTION

A typical gut microbiome consists of several hundred bacterial strains that span at least six orders of magnitude in relative abundance. Determining the contribution of individual strains to community ecology and host physiology is a daunting challenge. A variety of studies have characterized the functional properties of bacterial strains from the gut microbiome (Buffie et al., 2014; Mahowald et al., 2009; Marion et al., 2019, 2020; Patnode et al., 2019; Rey et al., 2010; Ridlon et al., 2020; Samuel and Gordon, 2006; Streidl et al., 2021; Studer et al., 2016). However, in vivo demonstrations of function typically involve mice colonized by one species or a small community; it remains difficult to study the functional contribution of a strain in the context of a native-scale community.

In thinking about where to start, two considerations led to the same idea. First, we were concerned about functional redundancy (Louca et al., 2018; Tian et al., 2020). If we drop out a single strain, will we fail to see a phenotype because a strain with a similar function is present in the community? Second, the intestinal ecosystem is organized into physical and metabolic niches, which are thought to serve as functional units within the community (Louca et al., 2018; Tian et al., 2020). A long-standing set of questions concerns how strains function within a niche. What is the mapping of strains to niches? Can changes in one niche can propagate to others? And how do these events govern emergent behaviors such as community architecture and metabolic output?

Taking these considerations into account, we decided to interrogate a niche rather than an individual strain. Given our interest in the chemistry of the microbiome, we focused on the metabolic niche surrounding a well-studied microbial pathway—bile acid 7α-dehydroxylation—reasoning that it generates a highly concentrated pool of metabolites with important biological activities (Funabashi et al., 2020; Ridlon et al., 2006, 2016), but its dynamics are poorly understood.

We took advantage of a recently developed model system for the gut microbiome that is composed of >100 of the most common gut bacterial species (Cheng et al., 2021) (**Table S1**). We find that the niche consists of two strains, *Clostridium scindens* (*Cs*) and *Clostridium hylemonae* (*Ch*); when we drop them out of the community together (Δ*Cs*Δ*Ch*), eight strains go up or down sharply in relative abundance. Single-strain dropout communities (Δ*Cs* and Δ*Ch*) reveal that either strain alone can metabolize bile acids exhaustively, and compensation within the niche keeps the relative abundance of its members within a narrow range. They also enable us to test causality in each strain-strain interaction, establishing whether it is specific to *Cs* or *Ch* or a function of both strains. Δ*Cs*- and Δ*Ch*-colonized mice have similar bile acid profiles but a large and unexpected difference in phenylalanine metabolism owing to a *Cs-*specific interaction with *Clostridium sporogenes*, showing that a strain swap within a niche can have a cascading effect that impacts unrelated niches. Finally, we show that *Cs* is sufficient to promote steatosis in mice fed a high-fat diet. These data show that a highly controlled experiment can be performed on a complex community to elucidate strain-level causality and mechanism.

### A computational search to identify strains that carry out bile acid 7α-dehydroxylation

We began by determining which strains in our 107-member community (hereafter, hCom1a) occupy the 7α-dehydroxylation niche. To address this question, we performed a multigeneblast search against a database of genome sequences from hCom1a for the eight-gene *bai* operon (**Figure 1A**), which encodes a metabolic pathway for the 7α-dehydroxylation of cholic acid (CA) and chenodeoxycholic acid (CDCA) to deoxycholic acid (DCA) and lithocholic acid (LCA), respectively (Funabashi et al., 2020) (**Figure 1B**). Only two of the 107 strains harbor the *bai* operon: *Clostridium scindens* ATCC 35704 (*Cs*) and *Clostridium hylemonae* DSM 15053 (*Ch*), both of which are known to generate secondary bile acids (Ridlon et al., 2016).

**Figure 1:**
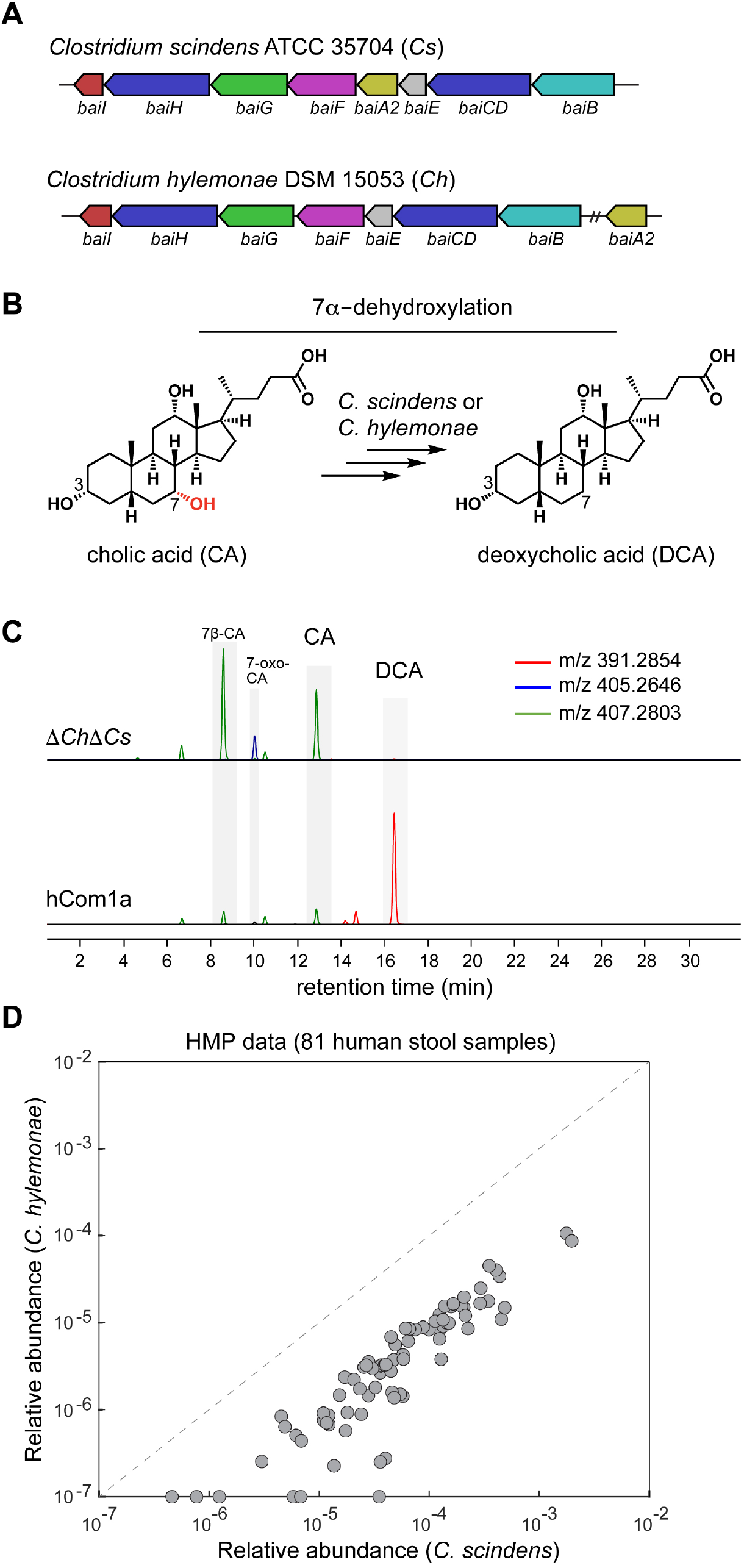
The 7α-dehydroxylation niche in hCom1a is composed of *Clostridium scindens* (*Cs*) and *Clostridium hylemonae* (*Ch*). (**A**) A multigeneblast search of the 107 genomes in hCom1a shows that only *Clostridium scindens* and *Clostridium hylemonae* harbor the *bai* operon, which encodes the bile acid 7α-dehydroxylation pathway. (**B**) A simplified schematic showing the dehydroxylation of cholic acid (CA) to deoxycholic acid (DCA). (**C**) Combined extracted ion chromatogram showing that hCom1a converts CA to DCA in vitro, whereas the two-strain dropout community Δ*Ch*Δ*Cs* does not. Constituent strains were cultured separately, pooled, and subcultured (1:100) in Mega medium containing 100 µM cholic acid for 72 h. Culture supernatants were collected and analyzed by LC-MS. (**D**) *Clostridium scindens* and *Clostridium hylemonae* typically co-colonize the human gut, as shown by their relative abundances in 81 healthy human stool samples from the NIH HMP.

Next, we sought to test whether any additional strains in the community harbor an alternative (unknown) pathway. We constructed hCom1a and a Δ*Ch*Δ*Cs* dropout community by mixing individually cultured strains, and then we grew these communities in the presence of CA and profiled their culture supernatants by LC-MS. We find that hCom1a converts CA to DCA whereas Δ*Ch*Δ*Cs* does not (**Figure 1C**). These data suggest that *Cs* and *Ch* are the only strains in the 7α-dehydroxylation niche, but they do not exclude the possibility that another strain is capable of carrying out this reaction in vivo. By analyzing publicly available human metagenomic data (Kraal et al., 2014), we observe that *Cs* and *Ch* often co-exist in the gut community, indicating that they might share this niche in the human gut (**Figure 1D, Table S1**).

### Determining the occupancy of the 7α-dehydroxylation niche in vivo

To test whether *Cs* and *Ch* are the only occupants of the 7α-dehydroxylation niche, we colonized germ-free C57BL/6 mice with hCom1a or the two-strain dropout community (Δ*Ch*Δ*Cs*). After three weeks, we sacrificed the mice, harvested intestinal contents, and subjected the samples to targeted metabolomic and high-resolution metagenomic analysis (**Figure 2A**). By analyzing metagenomic data, we verified the absence of *Cs* and *Ch* in cecal samples from the Δ*Ch*Δ*Cs*-colonized mice (**Figure 2B** and **S2A, Table S2**).

**Figure 2:**
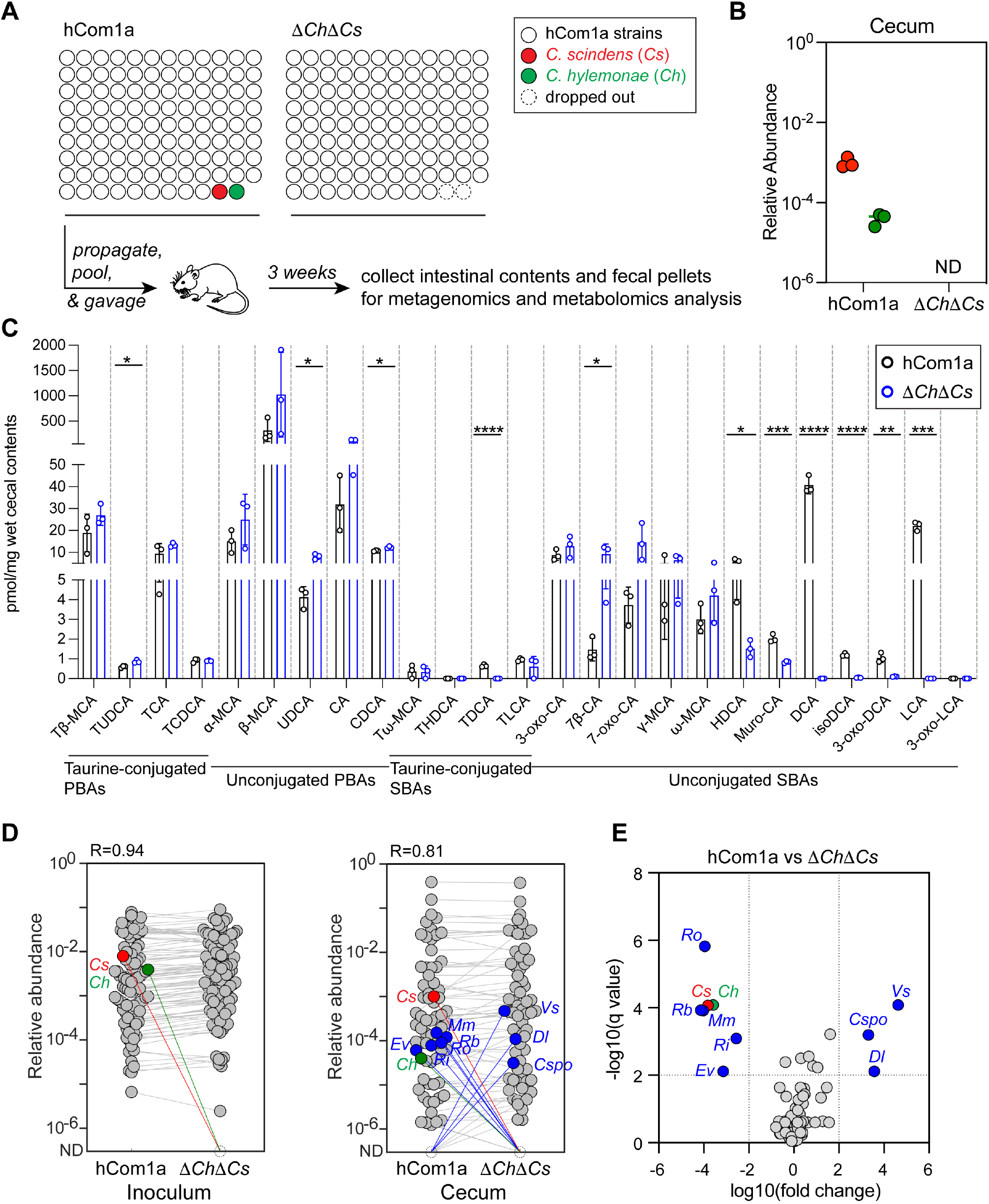
Metabolic and ecological impacts of removing strains in the 7α-dehydroxylation niche. (**A**) Schematic of the experiment. Germ-free C57BL/6 mice were colonized with hCom1a or Δ*Ch*Δ*Cs* and housed for 3 weeks before sacrifice. Fecal pellets and intestinal contents were subjected to metagenomic analysis or targeted metabolite profiling. (**B**) *Cs* and *Ch* are undetectable in Δ*Ch*Δ*Cs*-colonized mice. Relative abundances were calculated through a high-resolution metagenomic analysis of the inoculum and cecal communities. (**C**) Secondary bile acids are eliminated in Δ*Ch*Δ*Cs*-colonized mice. Bile acids were quantified in cecal contents by targeted LC-MS-based profiling. Statistical significance was assessed using a Student’s two-tailed t-test (*: p<0.05; **: p<0.01; ***: p<0.001; ****: p<0.0001). PBAs: primary bile acids; SBAs: secondary bile acids. (**D**) Average relative abundances of the inoculum (left) versus the cecal communities at week 3 (right). Each dot is an individual strain; the collection of dots in a column represents the community averaged over 3 mice co-housed in one cage. *Cs* and *Ch* are highlighted in red and green, respectively. Strains highlighted in blue went up or down in relative abundance between hCom1a-colonized and Δ*Ch*Δ*Cs*-colonized mice (FDR < 0.01, fold change > 100). (**E**) Volcano plot of differential strain relative abundance. The log10(fold change) values of each strain are shown; relative abundances were set at 10^−8^ for strains not detected. Relative abundances were analyzed using a multiple unpaired t test, corrected with FDR (Q<1%). Strains discovered with significantly different relative abundance (FDR<0.01, fold change >100) are colored blue; the full names of the blue strains can be found in **Figures 4D** and **S2**.

The removal of *Ch* and *Cs* has a large impact on the bile acid pool (**Figure 2C, Table S3**). The products of 7α-dehydroxylation—the secondary bile acids DCA and LCA—are absent in fecal pellets from the Δ*Ch*Δ*Cs* mice, as are prominent derivatives including isoDCA and 3-oxo-DCA. In contrast, Δ*Ch*Δ*Cs* mice have a higher level of the pathway intermediates 7β-CA and 7-oxo-CA and the CDCA epimer ursodeoxycholic acid (UDCA) (**Figure 2C**). Together, these data suggest *Cs* and *Ch* are the only strains in the 7α-dehydroxylation niche and that dropping them out results in the complete elimination of secondary bile acids. Previous attempts to knock out secondary bile acid production were conducted in the context of simple defined communities (Marion et al., 2019, 2020; Streidl et al., 2021; Studer et al., 2016) or by antibiotic knockdown of resident colonists followed by colonization with an antibiotic-resistant mutant, which depletes a variety of other strains in the community (Jin et al., 2022). In contrast, our system is a defined knockout of 7α-dehydroxylation in the setting of a complex community.

### Eliminating *Cs* and *Ch* has a large impact on a subset of strains

Next, we sought to assess the ecological impact of *Cs*/*Ch* removal on the rest of the community. Removing a species from a natural ecosystem can have consequences that are difficult to predict (Díaz et al., 2003; Wootton, 2010). In mice, species have been removed from simple defined communities (Marion et al., 2019, 2020; Streidl et al., 2021; Studer et al., 2016) or from undefined communities using antibiotic pretreatment and phage (Lam et al., 2021), but it remains unclear what effect species removal will have on a complex, unperturbed community in its native setting.

hCom1a and Δ*Ch*Δ*Cs* assume a broadly similar architecture in mice (R=0.84) (**Figure 2D, Table S2**). While 97/107 of the community members have a similar relative abundance, the removal of *Cs* and *Ch* has a striking effect on a subset of eight strains, using a strict threshold of q<0.01 and fold change >100 (**Figure 2D-E**). *Veillonella sp*. 3-1-44 (*Vs*), *Clostridium sporogenes* (*Cspo*), and *Dorea longicatena* (*Dl*) went up sharply in relative abundance, while *Ruminococcus obeum* (*Ro*), *Ruminococcus bromii* (*Rb*), *Mitsuokella multacida* (*Mm*), *Roseburia intestinalis* (*Ri*), and *Eubacterium ventriosum* (*Ev*) decreased (**Figure 2D** and **S2B**). Thus, at least in this case, the removal of two strains from a complex gut community has a large effect on a confined subset of strains that lie outside the 7α-dehydroxylation niche.

### Compensation and functional redundancy within the 7α-dehydroxylation niche

To gain more resolution into dynamics within the 7α-dehydroxylation niche, we constructed two new communities*—*Δ*Ch* and Δ*Cs—*in which the 7α-dehydroxylation niche is occupied by *Cs* or *Ch*, but not both. We colonized germ-free C57BL/6 mice with each of these communities or the parental community (hCom1a) and sampled fecal pellets weekly for three weeks. We then sacrificed the mice, harvested intestinal contents, and subjected all the samples to bile acid profiling and metagenomic sequencing with high-resolution read mapping (**Figure 3A**).

**Figure 3:**
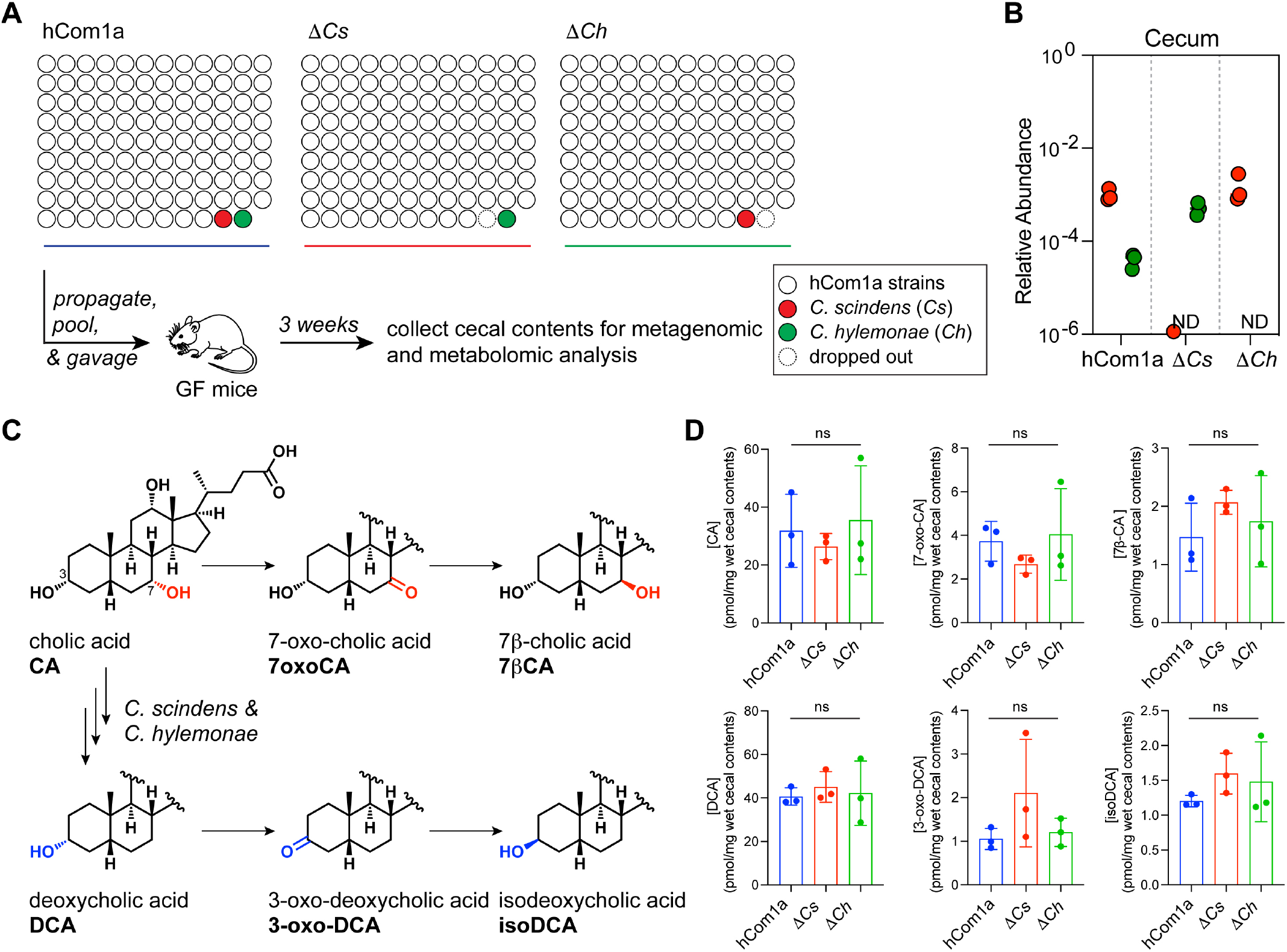
Compensation and functional redundancy within the 7α-dehydroxylation niche. (**A**) Schematic of the experiment. Germ-free C57BL/6 mice were colonized with hCom1a, Δ*Cs*, or Δ*Ch* and housed for 3 weeks before sacrifice. Cecal contents were subjected to metagenomic and targeted metabolomic profiling. (**B**) Compensation within the 7α-dehydroxylation niche keeps the total relative abundance of its residents similar. Relative abundances of *Cs* and *Ch* in cecal contents from hCom1a-, Δ*Ch*-, and Δ*Cs*-colonized mice are shown. (**C**) Metabolic pathways for bile acid transformation by the gut microbiota. (**D**) The bile acid pools of mice colonized by hCom1a, Δ*Ch*, and Δ*Cs* are comparable. Thus, in the context of a complete community, *Ch* and *Cs* can carry out the core function of the niche—the conversion of primary to secondary bile acids—on its own.

First, we validated that *Ch* and *Cs* were absent from their respective dropout communities in the cecal samples from week 3 (**Figure 3B** and **S2A, Table S2**). Next, we analyzed dynamics within the niche by metagenomic sequence analysis. In hCom1a, the relative abundances of *Cs* and *Ch* are ~10^−3^ and ~10^−4^, respectively. Consistent with the human data (**Figure 1D**), *Cs* is present at a higher relative abundance than *Ch* when they co-occupy the niche. In the absence of *Cs, Ch* goes up 12-fold in relative abundance; likewise, but to a lesser extent, *Cs* is more abundant in the Δ*Ch* community (**Figure 3B**). Thus, *Cs* and *Ch* can co-exist, but compensation within the niche keeps the total relative abundance of its residents at a similar level (1.0 × 10^−3^ in hCom1a, 5.1 × 10^−4^ in Δ*Cs*, and 1.5 × 10^−3^ in Δ*Ch*) (**Figure S2C**).

Finally, we quantified the cecal bile acid pool by LC-MS (**Figures 3C** and **S3A, Table S3**). The bile acid profiles of Δ*Ch*, Δ*Cs*, and the parental (hCom1a) community are remarkably similar: CA, DCA, and their derivatives (7-oxoCA, 7β-CA, 3-oxoDCA, and isoDCA) are all produced at comparable levels (**Figure 3D** and **S3A**). Thus, in the context of a complete community, either strain can carry out the core function of the niche—the conversion of primary to secondary bile acids—on its own.

### Single-strain dropouts reveal complex interactions among *Cs, Ch*, and interacting strains

Next, we turned to effects outside the 7α-dehydroxylation niche. To test whether the strain-level interactions observed in the Δ*Ch*Δ*Cs* community are caused by *Cs* or *Ch*, we compared metagenomic data from hCom1a-colonized mice with that of Δ*Cs-* and Δ*Ch*-colonized mice to assess the relative abundances of the rest of the strains in the community (**Figure 4A, Table S2**). The architectures of Δ*Cs* and Δ*Ch* are very similar to that of hCom1a (R=0.86 and 0.88, respectively) (**Figure 4B-C**). Interestingly, of the eight strains whose relative abundances increased or decreased by >100-fold in the Δ*Ch*Δ*Cs* community, five respond to the absence of *Cs* (*Vs* and *Cspo* ↑; *Ro, Rb*, and *Mm* ↓) and three are impacted by the absence of *Ch* (*Vs* ↑; *Ri* and *Mm* ↓) (**Figure 4B-C** and **S2B**). These sets overlap partially, suggesting that certain interactions share a common mechanism while others are *Cs-* or *Ch-*specific.

**Figure 4:**
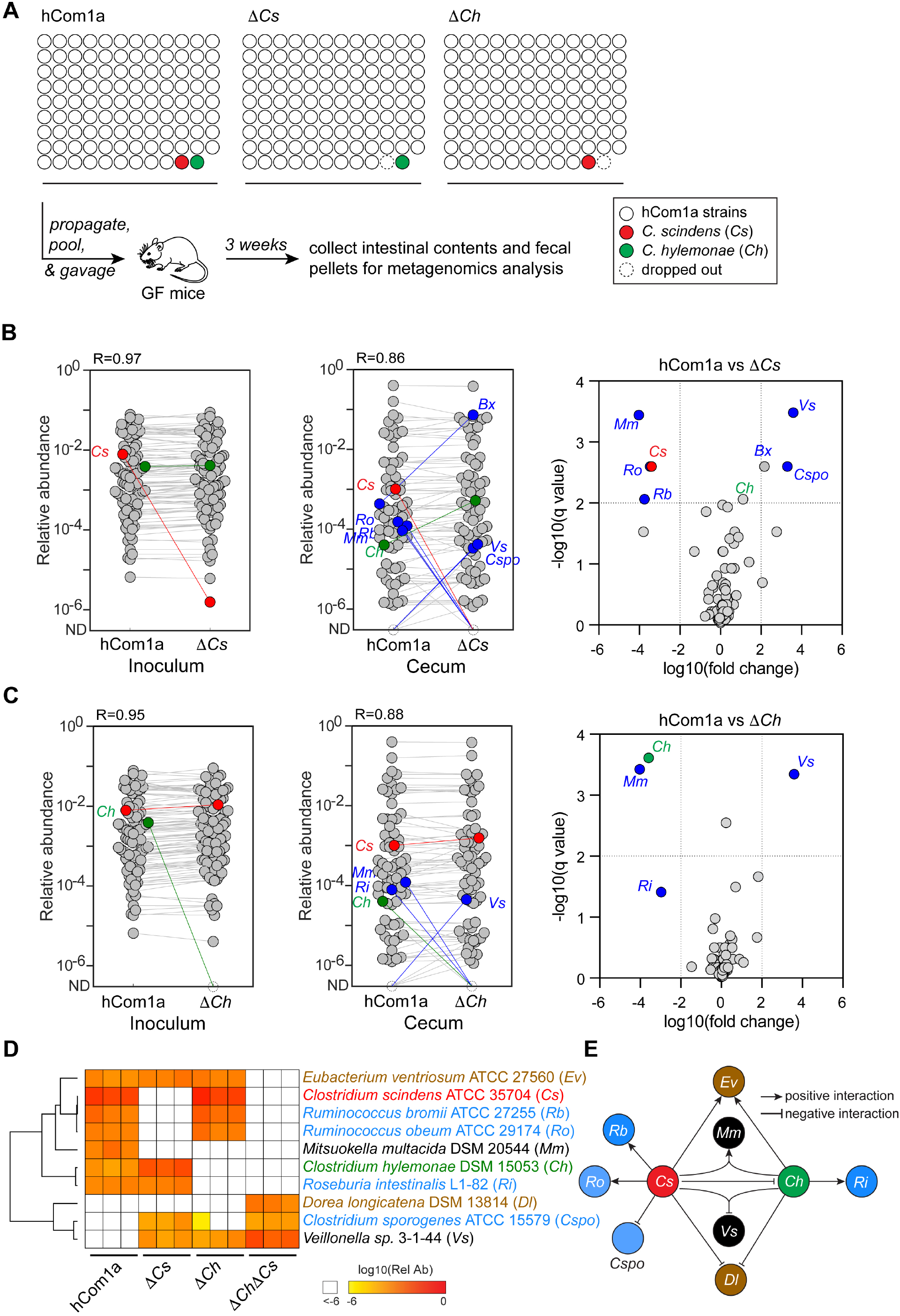
Single-strain dropouts reveal complex interactions among *Cs, Ch*, and interacting strains. (**A**) Schematic of the experiment. Germ-free C57BL/6 mice were colonized with hCom1a, Δ*Cs*, or Δ*Ch* and housed for 3 weeks before sacrifice. Fecal pellets and intestinal contents were subjected to metagenomic analysis. (**B-C**) While most strains remain unchanged when dropping out *Cs* or *Ch*, a small number of strains change in relative abundance >100-fold. *Left:* Metagenomic analysis showing that *Cs* and *Ch* are absent in the Δ*Cs* and Δ*Ch* community inocula, respectively. *Middle:* Dropping out *Cs* or *Ch* impacts the relative abundance of five or three strains in the Δ*Cs* and Δ*Ch* communities, respectively. Each dot is an individual strain; the collection of dots in a column represents the community averaged over three mice co-housed in one cage. *Cs* and *Ch* are highlighted in red and green. Strains colored blue went up or down in relative abundance between hCom1a-colonized and Δ*Cs* or Δ*Ch*-colonized mice (FDR<0.01, fold change >100). *Right:* Volcano plot showing the log10(fold change) values for each strain; for strains that were not detected, relative abundances were set at 10^−8^. Strains with significantly different relative abundance (FDR <0.01, fold change >10^2^) are colored blue; the full names of these strains are shown next to the heatmap in (**D**). (**D**) Heatmap representing the 8-strain interaction network around *Cs* and *Ch*. The relative abundance of each strain is shown in three mice colonized by hCom1a, Δ*Cs*, Δ*Ch*, or Δ*Ch*Δ*Cs*. Strains with relative abundance <10^−6^ are colored white. (**E**) Schematic of the interaction network. Strains are linked to the 7α-dehydroxylation niche in one of three ways: 1) Some strains are specific to *Cs* (*Rb, Ro*, and *Cspo*) or *Ch* (*Ri*); these interactions are presumably strain-specific and unrelated to bile acids. 2) *Ev* and *Dl* only respond to the double-strain dropout, indicating a mechanism related to secondary bile acid production. 3) The remaining strains, *Mm* and *Vs*, respond when either or both strains are missing, suggesting a requirement for the simultaneous presence of *Cs* and *Ch*.

To explore the interaction network around the 7α-dehydroxylation niche in more detail, we combined and clustered the metagenomics data from *Cs, Ch*, and each of the 8 affected strains (**Figure 4D**). The data are consistent with a model in which strains are linked to the niche in one of three ways: 1) Some strains are specific to *Cs* (*Rb, Ro*, and *Cspo*) or *Ch* (*Ri*); these interactions are presumably strain-specific and unrelated to bile acids. 2) *Ev* and *Dl* only respond to the double-strain dropout, indicating a mechanism related to secondary bile acid production. 3) The remaining strains, *Mm* and *Vs*, respond when either or both strains are missing, suggesting a requirement for the simultaneous presence of *Cs* and *Ch* (**Figure 4E**). We note that we cannot distinguish between direct interactions with *Cs* and/or *Ch* and indirect interactions that involve a third strain. Nevertheless, this analysis demonstrates the power of single-strain dropouts in discovering strain-strain interactions in a complex community.

### *Dl* growth is inhibited by a product of 7α-dehydroxylation

Next, we sought to gain insight into the mechanisms that govern the negative (inhibitory) interactions revealed by this analysis. We focused on *Dorea longicatena* (*Dl*) and *Clostridium sporogenes* (*Cspo*) since *Veillonella sp*. 3-1-44 (*Vs*) did not grow robustly in vitro. *Dl* is only detectable in vivo when *Cs* and *Ch* are both absent, so we reasoned that either *Cs* or *Ch* should be capable of inhibiting its growth. We hypothesized that this growth inhibition is mediated by secondary bile acids, which are produced by both strains. To test this hypothesis, we cultured *Dl* (and, as a control, *Cspo*) in growth medium supplemented with CA and DCA at concentrations ranging from 39 µM-1.25 mM; CDCA and LCA were not considered due to poor solubility at high concentrations. The growth of *Dl* is completely inhibited by DCA at 625 µM, a physiological concentration, while *Cspo* was only partially affected (**Figure S4A**). In contrast, neither strain was affected by CA at concentrations as high as 1.25 mM (**Figure S4B**). These results suggest that DCA, which is produced by *Cs* and *Ch*, inhibits colonization by *Dl*. Although *C. sporogenes* is moderately affected by DCA, other mechanisms are more likely responsible for its ability to bloom only when *Cs* is absent (Kang et al., 2019).

### An unexpected impact of *Cs* on aromatic amino acid metabolism

The analysis above suggests that although *Ch* and *Cs* appear functionally redundant within their niche—the two strains are interchangeable in terms of the resulting bile acid profile—they have distinct effects on the rest of the community. However, there are only six strains whose relative abundances are meaningfully different between the Δ*Ch* and Δ*Cs* communities (*Bx, Ri, Rb, Ro, Mm*, and *Cspo*); apart from these, the relative abundances of the rest of the community are nearly superimposable (**Figure S5A, Table S2**). To determine whether the Δ*Ch* and Δ*Cs* communities are truly indistinguishable, we compared microbiome-derived metabolites in the cecal contents and urine of the Δ*Ch*- and Δ*Cs*-colonized mice, expecting to find concordant profiles in light of the compositional similarities (**Figure 5A**).

**Figure 5:**
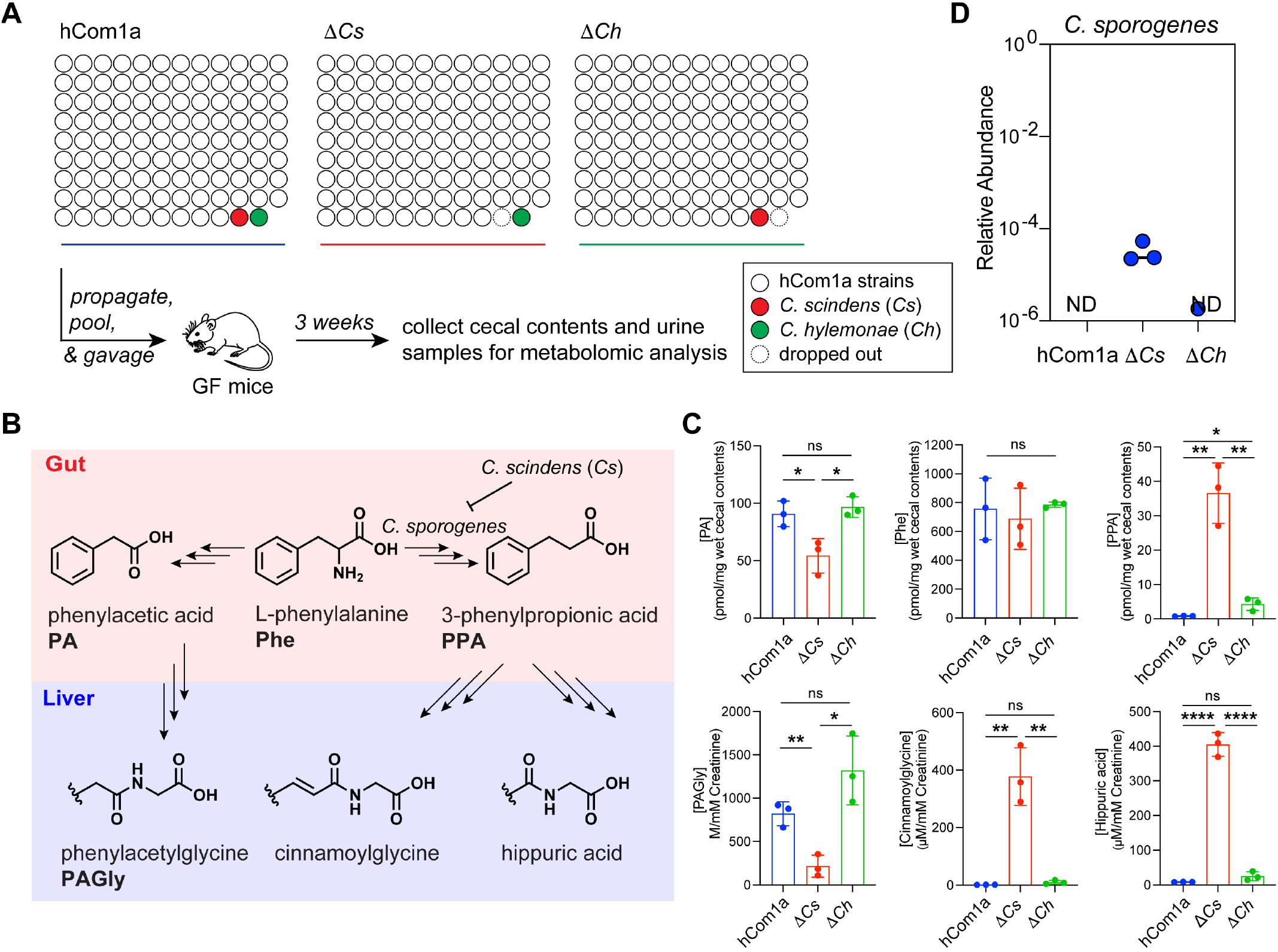
An unexpected impact of *Cs* on aromatic amino acid metabolism. (**A**) Schematic of the experiment. Germ-free C57BL/6 mice were colonized with hCom1a or its single-strain dropout variant (Δ*Cs* and Δ*Ch*) and housed for 3 weeks before sacrifice. Fecal pellets, cecal contents, and urine samples were subjected to targeted metabolite profiling. (**B**) *C. sporogenes*, which is undetectable in hCom1a- and Δ*Ch*-colonized mice, rises to a relative abundance of 10^−5^-10^−4^ in Δ*Cs*-colonized mice. The relative abundance of *Cspo* in cecal contents from hCom1a-, Δ*Cs*-, and Δ*Ch*-colonized mice is shown. (**C**) Certain gut bacteria can reduce phenylalanine (Phe) to phenylpropionic acid (PPA), which is converted to hippuric acid and cinnamoylglycine by the host. Other gut bacteria oxidize Phe to phenylacetic acid (PA), which is metabolized in the liver to phenylacetylglycine (PAGly). (**D**) Δ*Cs* differs markedly from hCom1a and Δ*Ch* in terms of AAA metabolite output. hCom1a and Δ*Ch* convert phenylalanine almost exclusively to PA (from which the host generates PAGly); hippurate is nearly undetectable. In contrast. Δ*Cs* converts Phe predominantly to phenylpropionic acid (which the host metabolizes to hippurate); PAGly levels are very low. Statistical significance was assessed using a Student’s two tailed t-test (*: p<0.05; **: p<0.01; ***: p<0.001; ****: p<0.0001, n.s: no significance).

To our surprise, there were striking differences among the profiles (**Figure 5B, Table S3**). First, three reductive phenylalanine metabolites—phenylpropionic acid, hippuric acid, and cinnamoylglycine— were produced abundantly in Δ*Cs*-colonized mice, but at very low levels in Δ*Ch*- and hCom1a-colonized mice (**Figure 5C** and **S3B-C**). Two observations suggest a causal role for *Cspo:* it is the only strain that is essentially undetectable in Δ*Ch*- and hCom1a-colonized mice but up >1000-fold in Δ*Cs*-colonized mice (**Figure 5D**), and it is known to produce reductive phenylalanine metabolites in vitro and in vivo (Dodd et al., 2017; Han et al., 2021).

Second, two oxidative metabolites of phenylalanine—phenylacetic acid and phenylacetylglycine— show the opposite pattern (**Figure 5C** and **S3B-C**). Given that reductive and oxidative pathways are competing since they share phenylalanine as a substrate (Dodd et al., 2017), the decrease in oxidative metabolites likely results from the increase in *C. sporogenes*-mediated reductive metabolism.

These data are consistent with a model in which a strain swap (Δ*Ch vs* Δ*Cs*) initiated a multi-step process that cascaded through the community: *Cs* was replaced with *Ch*, which increased the relative abundance of *Cspo* from nearly undetectable to ~10^−4^, which increased the level of hippurate (~16 fold) and cinnamoylglycine (~37 fold), which decreased the production of phenylacetate (~2 fold) and phenylacetylglycine (~6 fold) (**Figure S5B**).

This is especially notable in light of the biological activities of the metabolites involved. Phenylacetylglycine, which plays a causative role in cardiovascular disease (Nemet et al., 2020), is down substantially in Δ*Cs*-colonized mice. It is replaced by hippuric acid and cinnamoylglycine, which are notable for their lack of toxicity (Hoskins et al., 1984; Lees et al., 2013). Replacing *Cs* with *Ch* results in a 133-fold difference in the ratio of favorable (hippurate and cinnamoylglycine) to unfavorable (phenylacetylglycine) Phe metabolites. Thus, a strain swap in the 7α-dehydroxylation niche is ‘silent’ in terms of bile acids but yields a large, desirable effect in an unrelated compartment of the community.

### Constructing a *Cs* dropout in high-fat-fed mice

Finally, we returned to one of the questions that motivated this work: do individual strains from the gut community make a measurable contribution to host physiology? Specifically, we sought to test the hypothesis that a single strain capable of 7α-dehydroxylation can impact physiology via the production of secondary bile acids.

Having shown that the Δ*Ch*Δ*Cs* community is null for secondary bile acids while the Δ*Ch* community produces wild-type levels, we sought to compare the two communities, which differ only the presence of *Cs*. We colonized germ-free C57BL/6 mice with both communities and fed the mice standard chow for two weeks. We then switched to the Gubra Amylin NASH (GAN) diet, which is rich in saturated fat (40%) and commonly used to model fibrosing NASH (Hansen et al., 2020), and housed mice for another eight weeks (**Figure 6A**).

**Figure 6:**
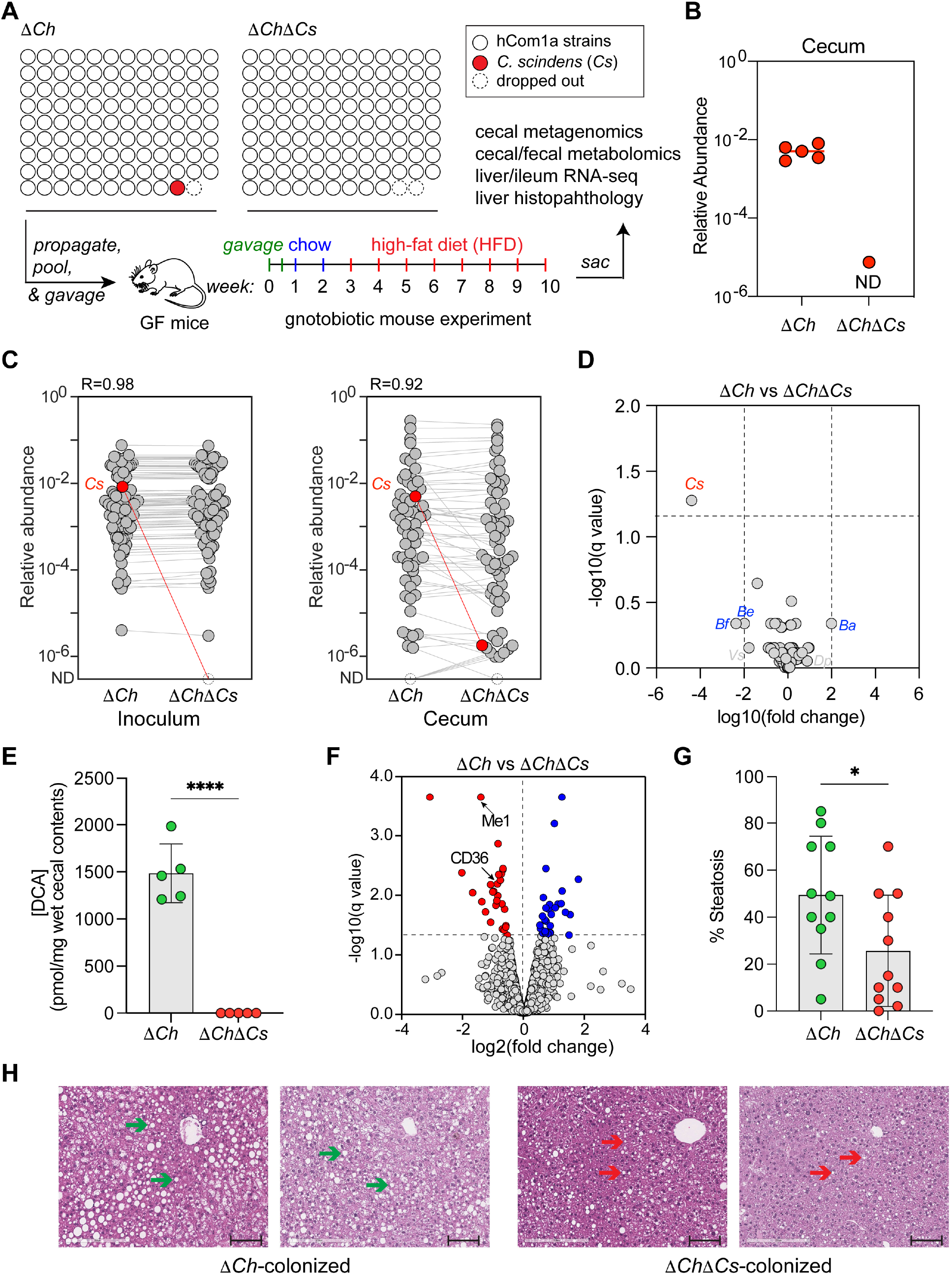
*Cs* dropout reveals a causal role in promoting steatosis. (**A**) Schematic of the experiment. Germ-free C57BL/6 mice were colonized with Δ*Ch* or Δ*Ch*Δ*Cs* and house for 10 weeks before sacrifice. Mice were fed chow for two weeks and then a high-fat diet (HFD) for 8 weeks. Fecal pellets and cecal contents were subjected to metabolomics and targeted metabolite profiling, while liver and ileal tissue were analyzed by RNA-seq. Liver sections were stained with H&E and assessed for steatosis. (**B** and **C**) Metagenomic analysis showing that the indicated strains are absent in the Δ*Ch* and Δ*Ch*Δ*Cs* inocula and cecal contents, respectively. (**D**) The composition of the Δ*Ch* and Δ*Ch*Δ*Cs* communities in mice fed a HFD is very similar. The volcano plot shows the log10(relative abundance) values for each strain; for strains that were not detected, relative abundances were set at 10^−8^. Strains with significantly different relative abundance (fold change >10^2^) are colored blue. (**E**) Dropping out *Cs* from the Δ*Ch* community eliminates the production of the 7α-dehydroxylation pathway product DCA. Bile acids were quantified in cecal contents using LC-MS. (**F**) Transcriptomic analysis showing host genes that are differentially expressed in the presence and absence of *Cs*. Genes whose expression level changed significantly (q<0.05) are highlighted in blue (up-regulated) or red (down-regulated). (**G**) The extent of steatosis in H&E-stained liver sections was assessed by a board-certified pathologist in a blinded manner. (**H**) Representative H&E-stained liver sections from Δ*Ch* or Δ*Ch*Δ*Cs*-colonized mice fed with a HFD in two independent experiments. Scale bars, 100 μm. Green and red arrows indicate representative lipid droplets within hepatocytes. The data in (**B**-**F**) are from a representative experiment with one technical replicate (inoculum) or five biological replicates (cecum). Data in (**G** and **H**) come from two independent experiments; n = 11 mice per group total. In (**E**) and (**G**), data were analyzed using unpaired two-tailed Student’s t test, and the asterisk indicates p<0.05 (*) or p<0.0001(****).

First, we analyzed metagenomic sequencing data from the inoculum and from cecal contents at the time of sacrifice to validate the dropout and assess community composition in mice fed a high-fat diet (**Table S4**). *Cs* was present in the inoculum and cecal contents of the Δ*Ch* community but absent in Δ*Ch*Δ*Cs*, confirming that the communities had the intended composition (**Figure 6B**). The relative abundances of other community members were broadly similar between the two groups of mice (R=0.92) (**Figure 6C**). Impacts on other strains were blunted in mice fed a HFD (**Figure 6D**). Taken together, these data show that the Δ*Ch* and Δ*Ch*Δ*Cs* communities have a comparable architecture in vivo under conditions of high-fat feeding.

To characterize the effect of *Cs* on the bile acid pool in mice fed a high-fat diet, we performed targeted metabolite profiling on the cecal contents (**Table S4**). As before, we observed that DCA, LCA, isoDCA, and 3-oxo-DCA are present in Δ*Ch*-colonized mice and absent in Δ*Ch*Δ*Cs*-colonized mice (**Figures 6E** and **S6**). Δ*Ch*Δ*Cs*-colonized mice have a higher level of the pathway substrates CA and CDCA, their derivatives 7β-CA and 7-oxo-CA, and the CDCA epimer UDCA (**Figure S6**). The taurine-conjugated primary bile acids TCA and TMCA are produced at a comparable level in the two communities, indicating that bile salt hydrolase activity was not significantly affected (**Figure S6**). Taken together, these data suggest that *Cs* alone is sufficient to restore 7α-dehydroxylation in vivo.

Notably, the bile acid pool in mice fed a high-fat diet is much larger than mice fed conventional chow. For example, the level of DCA in Δ*Ch*-colonized mice is >30-fold higher than that in the chow condition (1486 vs 42 pmol/mg wet cecal contents), which is consistent with a 5-fold increase in the relative abundance of *Cs* in Δ*Ch*-colonized mice in mice fed a HFD (5 × 10^−3^) vs. mice fed conventional chow (1 × 10^−3^).

### *Cs* dropout reveals a causal role in promoting steatosis

As a starting point for assessing the impact of *Cs* on host physiology, we measured the impact of *Cs* dropout on the host transcriptome. We performed RNA sequencing (RNA-seq) on the small intestine and liver of Δ*Ch*- and Δ*Ch*Δ*Cs*-colonized mice, sites of bile acid residence during enterohepatic circulation (De Aguiar Vallim et al., 2013; Wahlström et al., 2016). Only 67 genes were differentially expressed in the liver and 22 in the ileum (q<0.05) (**Figure 6F** and **S7A-C, Table S4**).

Two genes down-regulated in the liver of Δ*Ch*Δ*Cs*-colonized mice—*CD36* and *Me1*—caught our attention due to their function in lipid metabolism. CD36 (fatty acid translocase, FAT) plays a pathogenic role in hepatic steatosis in mice by facilitating the uptake and intracellular transport of long-chain fatty acids (Koonen et al., 2007; Wilson et al., 2016); clinical studies have reinforced its significance by showing increased expression in the liver of NAFLD patients (Rada et al., 2020). Me1 (cytosolic malic enzyme) catalyzes the oxidative decarboxylation of malate to pyruvate, CO_2_ and NADPH, contributing to de novo fatty acid synthesis (Wise and Ball, 1964) and hepatic steatosis (Al-Dwairi et al., 2012). The difference in transcript and protein levels of CD36 and Me1 was confirmed by RT-qPCR and western blot (**Figure S7D-E, Table S4**).

In light of these results, we hypothesized that secondary bile acids produced by *Cs* would impact lipid absorption, transport, and synthesis, especially in mice fed a high-fat diet. To test this hypothesis, we performed a blinded histologic assessment of H&E-stained liver slides to assess the extent of steatosis in Δ*Ch*- and Δ*Ch*Δ*Cs*-colonized mice at the time of sacrifice. We found that Δ*Ch*Δ*Cs*-colonized mice had a lower level of steatosis than Δ*Ch*-colonized mice, consistent with a model in which secondary bile acids contribute to fat deposition in the liver (**Figure 6G-H**). These data show that a single strain from the gut community can impact hepatic physiology, likely by means of a diffusible metabolite.

## DISCUSSION

A long-standing challenge in microbiome research has been to assess the impact of individual strains on community ecology and host physiology. Previous efforts have focused on interactions in binary culture (Medlock et al., 2018; Venturelli et al., 2018) or in simple communities (Gutiérrez and Garrido, 2019; Sanchez-Gorostiaga et al., 2019), where the rules and selective conditions are likely distinct from those in a complex community (Bairey et al., 2016). A complex defined community offers a setting in which a ‘clean’ dropout can be constructed in a physiologically relevant background. This approach, in combination with recent advances in genetic systems for the microbiome (García-Bayona and Comstock, 2019; Guo et al., 2019; Jin et al., 2022), will greatly improve our understanding of strain-strain and strain-host interactions.

Our results emphasize the importance of focusing on a functional unit within the community—here, a metabolic niche—rather than on individual strains. Initially, our main focus was to characterize the effects of secondary bile acids on the host, and our interest in the niche was driven mainly by the concern that the phenotype induced by dropping out a single bile-acid-producing colonist would be masked by another functionally redundant strain. But as we began analyzing metagenomic and metabolomic data from the strain dropout experiments, we came to think that the main story was the connection between ecological dynamics and community metabolism.

Using this model, we show that there is functional redundancy within the 7α-dehydroxylation niche. An unknown mechanism—we speculate that it could be the availability of the primary bile acids CA and CDCA—keeps the total relative abundance of strains in the niche within a 3-fold range, regardless of whether the niche is mono- or bi-colonized. *Cs* and *Ch* can co-occupy the niche rather than excluding each other. However, when the niche is mono-colonized, *Cs* and *Ch* are functionally interchangeable; either strain alone is capable of dehydroxylating the entire pool of primary bile acids, and the resulting bile acid profiles are indistinguishable.

However, ‘silent’ changes within the niche lead to unexpected effects elsewhere in the community. Communities in which the niche is mono-colonized with *Cs* (Δ*Ch*) vs. *Ch* (Δ*Cs*) are extremely similar in terms of architecture—only six strains differ significantly in relative abundance. But in Δ*Cs*, the unexpected proliferation of *Clostridium sporogenes*—which is below the limit of detection in hCom1a and Δ*Ch*—led to a large increase in reductive phenylalanine metabolites, reshaping the chemical output of a different niche in the community. This finding has two important implications. First, it redefines how we think about the functional impact of strain-level variation in the microbiome. Different strains that fill the same niche—even those that appear functionally redundant—may yield a large difference outside the niche. As a corollary, variation in a phenotype of interest may come from an unrelated place; if one were trying to figure out why hippurate levels were so low in hCom1a-colonized mice, it would not have been obvious to look in the 7α-dehydroxylation niche.

Second, this has important implications for understanding the principles of rational community design. An emerging goal in microbiome research is to design microbial communities that are endowed with an immunologic or metabolic phenotype of interest (Fischbach, 2018). If the goal is to make a community that generates hippurate and not phenylacetylglycine, the obvious strategy would be to include strains that produce the former and exclude those that generate the latter. But by dropping out a strain in an unrelated niche, we observed a comparable difference to what we would have expected through rational engineering—and through a simpler maneuver that does not reduce diversity or compromise any other niche. Thus, to alter a phenotype of interest, one needs to consider not just the strains that carry it out but their interaction partners in the community.

## MATERIALS AND METHODS

### Bacterial strains and culture conditions

All strains used in the synthetic community were obtained from American Type Culture Collection (ATCC), Leibniz Institute DSMZ-German Collection of Microorganisms and Cell Cultures GmbH (DSMZ), BEI resources (BEI), and other sources as indicated in **Table S1**. All strains were cultured in one of two growth media: mega medium (MM) and chopped meat medium w/ rumen fluid and carbohydrates (CMM) (**Table S1**). Cultures were incubated at 37 °C in an anaerobic chamber (Coy Laboratories) in an atmosphere of 5% hydrogen, 10% CO_2_ and 85% N_2_. Cultures were stored in anaerobically prepared 25% glycerol/water (v/v) in 12.7 × 49 mm cryogenic vials with closures (Corning 430659) or 1.2 ml V-bottom 96-well plates (E&K Scientific Products, Inc., OX1263-S) capped with a plate seal (E&K Scientific Products, Inc., EK-2066) and sealed with oxygen-impervious yellow vinyl tape (Coy Laboratories, 1600330w) to ensure anoxic conditions during long-term storage. All medium and reagents used in the anaerobic chamber were pre-reduced for at least 48 h.

### Synthetic community construction

Frozen stocks in a 96-well plate were thawed, and 100 μl of each thawed culture was used to inoculate 1 ml of growth medium (**Table S1**) in a sterile 2.2 ml 96-well plate (E&K Scientific Products, Inc., OX1265-S). Strains were sub-cultured by 1:10 dilution into fresh medium daily for 2-3 days; growth was monitored by optical density at 600 nm (OD_600_) using a microplate spectrophotometer (BioTeK, Epoch2). For each batch of inoculum, a few strains (typically <10) had a low OD (<0.1). For these strains, additional cells were harvested and added from a 10 ml liquid culture prepared from a glycerol stock, or from single colonies scraped from solid growth medium. Finally, non-normalized cultures of all strains were pooled into a mixture. A 1 ml aliquot of the resulting mixed culture was stored at −80 °C for metagenomic sequencing. The remainder of the mixed culture was subjected to centrifugation (5000 x *g*, 15 min); the cell pellet was washed with an equal volume of pre-reduced sterile phosphate-buffered saline (PBS), and then resuspended in 1/10 of the initial volume of a 25% glycerol/water (v/v) solution. 1.2 ml aliquots of the resulting synthetic community were stored in 2 ml cryovials (Corning, 430659) at −80 °C until use.

### Gnotobiotic mouse experiments

Germ free Swiss-Webster or C57BL/6 mice (male, 6-8 weeks of age) were originally obtained from Taconic Biosciences (Hudson, NY) or Charles River Laboratories (Wilmington, MA) and colonies were maintained in gnotobiotic isolators and fed ad libitum. The Institutional Animal Care and Use Committee (IACUC) at Stanford University or Cleveland Clinic approved all procedures involving animals.

Glycerol stocks of synthetic communities (~1.2 ml) were thawed and shaken well at room temperature, and mice were orally gavaged with ~200 µl of the mixed culture. To ensure efficient colonization by all strains in the community, mice were gavaged using the same procedure on three successive days for all experiments except the steatosis experiment (**Figure 6**), for which mice were gavaged twice in the first week.

For experiments in which mice were fed standard chow, fecal pellets were collected weekly and stored at −80 °C prior to analysis. The mice were maintained on a standard diet (LabDiet 5k67; 0.2% Trp) for 3-4 weeks before sacrifice. Mice were euthanized humanely by CO_2_ asphyxiation and the luminal contents of the small intestine, cecum, and colon were collected and stored at −80 °C until use.

For experiments involving a high-fat diet (HFD), germ-free male C57BL/6 mice were purchased from Charles River at 4 weeks of age and gavaged with synthetic communities as described above. Mice were maintained on LabDiet 5k67 (LabDiet, St. Louis, MO) for 2 weeks before changing to the HFD (Research Diets D09100310G) for another 8 weeks before sacrifice. Mice were maintained on a 14:10 light:dark cycle, received autoclaved non-acidified water, and were housed on ALPHA-dri Plus bedding (Shepherd Specialty Papers, Watertown TN, USA). Fecal pellets were collected biweekly and stored at −80 °C prior to analysis. The total body weight of each mouse was recorded weekly. Livers were harvested before fixing the right lobe in formalin and snap-freezing the left, median, and caudate lobes for subsequent analysis. The intestinal tract was then excised and separated into the duodenum, jejunum, ileum, cecum, and colon. The luminal contents of all intestinal segments were collected by flushing with 3 ml sterile 0.9% saline and collecting the contents in sterile tubes before snap freezing. The remaining intestinal tissue was snap-frozen for subsequent RNA-seq analysis.

### Metagenomic sequencing

The same experimental pipeline was used for sequencing bacterial isolates and microbial communities. Bacterial cells were pelleted by centrifugation in an anaerobic environment. Genomic DNA was extracted using the DNeasy PowerSoil HTP kit (Qiagen) and the quantity of extracted genomic DNA was measured in a 384-well format using the Quant-iT PicoGreen dsDNA Assay Kit (Thermo Fisher). Sequencing libraries were generated in a 384-well format using a custom, low-volume protocol based on the Nextera XT process (Illumina). Briefly, the DNA concentration from each sample was normalized to 0.18 ng/µl using a Mantis liquid handler (Formulatrix). In cases where the concentration was below 0.18 ng/µl, the sample was not diluted further. Tagmentation, neutralization, and PCR steps of the Nextera XT process were performed on the Mosquito HTS liquid handler (TTP Labtech), creating a final volume of 4 µl per library. During the PCR amplification step, custom 12-bp dual unique indices were introduced to eliminate barcode switching, a phenomenon that occurs on Illumina sequencing platforms with patterned flow cells (Sinha et al., 2017). Libraries were pooled at the desired relative molar ratios and cleaned up using Ampure XP beads (Beckman) to effect buffer removal and library size selection. The cleanup process was used to remove fragments shorter than 300 bp and longer than 1.5 kb. Final library pools were quality checked for size distribution and concentration using the Fragment Analyzer (Agilent) and qPCR (BioRad). Sequencing reads were generated using the NovaSeq S4 flow cell or the NextSeq High Output kit, both in 2×150 bp configuration. 5-10 million paired-end reads were targeted for bacterial isolates and 20–30 million paired end reads for bacterial communities.

### Metagenomic read mapping

Paired-end reads from each sample were aligned to the hCom1a database using Bowtie2 with maximum insert length (-maxins) set to 3000, maximum alignments (-k) set to 300, suppressed unpaired alignments (--no-mixed), suppressed discordant alignments (--no-discordant), suppressed output for unaligned reads (--no-unal), required global alignment (--end-to-end), and using the “--very-sensitive” alignment preset (command: --very-sensitive -maxinsX 3000 -k 300 --no-mixed --no-discordant --end-to-end --no-unal). The output was piped into Samtools v. 1.9 (Li et al. 2009), which was used to convert the alignment output from SAM output stream to BAM format and then sort and index the BAM file by coordinates. Alignments were filtered to only keep those with >99% identity for the entire length of the read.

The median percentage of unaligned reads was 4.95% (range 4.10% - 8.35%). To assess the origin of these reads, we performed a BLAST v2.11.0+ search through the ncbi/blast:latest docker image with parameters “-outfmt ‘6 std qlen slen qcovs sscinames staxids’ -dbsize 1000000, -num_alignments 100” from a representative sample against the ‘NCBI - nt’ database as on 2021-02-16. We then filtered the BLAST results to obtain the top hits for a given query. Briefly, the script defined top hits as ones that had an e-value <= 1e-30, percent identity >= 99% and were within 10 percent of the best bit score for that query. To visualize and summarize the output, we used the ktImportTaxonomy script from the Krona package with default parameters. Reads were aggregated by NCBI taxon id and separately by genus. We found that most of the hits are from taxa that are closely related to the organisms in our community, while others are from the mouse genome. We conclude that our experiments did not suffer from any appreciable level of contamination.

### Sample preparation for LC/MS

For mouse fecal samples (~40 mg) or cecal contents (~80 mg), wet tissues were pre-weighed into a 2 ml screw top tube containing six 6 mm ceramic beads (Precellys® CK28 Lysing Kit). 600 µL (for fecal samples) or 1 ml (for cecal contents) of a mixture of ice-cold acetonitrile, methanol, and water (4/4/2, v/v/v) was then added to each tube and samples were homogenized by vigorous shaking using a QIAGEN Tissue Lyser II at 25/s for 10 min. The resulting homogenates were subjected to centrifugation for 15 min at 4 °C at 18,000 x *g*. 100 μl of the supernatant was then combined with 100 µl of an aqueous solution of an internal standard (2 μM d^4^-cholic acid). The resulting mixtures were then filtered through a Durapore PVDF 0.22-μ m membrane using Ultrafree centrifugal filters (Millipore, UFC30GV00), or MultiScreen Solvinert 96 Well Filter Plate (Millipore, MSRLN0410), and 5 µl was injected into the LC/MS.

For mouse urine samples, 5 µl of a urine sample was diluted 10-fold with ddH2O and then mixed with 50 µl of an aqueous internal standard solution (20 μM 4-chloro-L-phenylalanine). After centrifugation for 15 min at 4 °C at 18,000 x *g*, 50 µl of the resulting mixture was used for quantification of creatinine using a Creatinine Assay Kit (ab204537) as described in the manufacturer’s protocol. The remaining 50 µl of each sample was filtered through a Durapore PVDF 0.22-μm membrane using Ultrafree centrifugal filters (Millipore, UFC30GV00), and 5 µl was injected into the LC/MS.

### Liquid chromatography/mass spectrometry (LC/MS)

#### Bile acids

compounds were separated using an Agilent 1290 Infinity II UPLC equipped with a Kinetex C18 column (1.7 µm, 2.1 × 100 mm, Phenomenex) and detected using an Agilent 6530 Q-TOF equipped with a dual Agilent jet stream electrospray ionization (AJS-ESI) source operating under extended dynamic range (EDR 1700 m/z) in negative ionization mode. The parameters of the AJS-ESI source were as follows: gas temp: 300 °C; drying gas: 7.0 l/min; nebulizer: 40 psig; sheath gas temp: 350 °C; sheath gas flow: 10.0 l/min; VCap: 3500 V; nozzle voltage: 1400V; and fragmenter: 200 V. Mobile phase A was 0.05% formic acid in H_2_O, and mobile phase B was 0.05% formic acid in acetone. 5 μl of each sample was injected via autosampler into mobile phase and chromatographic separation was carried out at a flow rate of 0.35 ml/min with a 32-min gradient condition (t = 0 min, 25% B; t = 1 min, 25% B; t = 25 min, 75% B, t = 26 min, 100% B, t = 30 min, 100% B, t = 32 min, 25% B).

#### Aromatic amino acid metabolites

compounds were separated using an Agilent 1290 Infinity II UPLC equipped with an ACQUITY UPLC BEH C18 column (1.7 μm, 2.1 mm x 150 mm, Waters) and detected using an Agilent 6530 Q-TOF equipped with a standard atmospheric-pressure chemical ionization (APCI) source or dual Agilent jet stream electrospray ionization (AJS-ESI) source operating under extended dynamic range (EDR 1700 m/z) in negative ionization mode. For the APCI source the parameters were as follows: gas temp: 350 °C; vaporizer: 350 °C; drying gas: 6.0 l/min; nebulizer: 60 psig; VCap: 3500 V; corona: 20 μA; and fragmenter: 135 V. For the AJS-ESI source the parameters were as follows: gas temp: 350 °C; drying gas: 10.0 l/min; nebulizer: 40 psig; sheath gas temp: 300 °C; sheath gas flow: 11.0 l/min; VCap: 3500 V; nozzle voltage: 1400V; and fragmenter: 130 V. Mobile phase A was 6.5 mM ammonium bicarbonate in H_2_O, and B was 6.5 mM ammonium bicarbonate in 95 % MeOH/H_2_O. 5 μl of each sample was injected via autosampler into mobile phase and chromatographic separation was carried out at a flow rate of 0.35 mL/min with a 10-min gradient condition (t = 0 min, 0.5% B; t = 4 min, 70% B; t = 4.5 min, 98% B; t = 5.4 min, 98% B; t = 5.6 min, 0.5% B).

Online mass calibration was performed using a second ionization source and a constant flow (5 μl/min) of reference solution (119.0363 and 966.0007 m/z). The MassHunter Quantitative Analysis Software (Agilent, version B.09.00) was used for peak integration based on retention time (tolerance of 0.2 min) and accurate m/z (tolerance of 30 ppm) of chemical standards. Quantification was based on a 2-fold dilution series of chemical standards spanning 0.098 to 200 μM (for AAA metabolites) or 0.001 to 200 μM (for bile acids) and measured amounts were normalized by weights of extracted tissue samples (pmol/mg wet tissue) or creatinine level in the urine sample (μM/mM creatinine). The linear quantification range and lower limit of detection for all metabolites are listed in **Table S1**. The MassHunter Qualitative Analysis Software (Agilent, version 7.0) was used for targeted feature extraction, allowing mass tolerances of 30 ppm.

### Bile acids inhibition assay

*D. longicatena* and *C. sporogenes* were streaked from a glycerol stock onto Columbia Agar with 5 % Sheep Blood (BD 221165) and incubated for ~ 24 h at 37 °C. Individual colonies were picked and used to inoculate 3 ml of Mega medium and cultured for 24 h at 37 °C. Cells were diluted 1,000-fold into fresh Mega medium supplemented with various concentration of CA or DCA (final concentration raging from 1.25 mM, 625 µM, 312 µM, 156 µM, 78 µM, to 39 µM) or DMSO blank control. The starting OD_600_ for each culture was 0.1. 200 µl aliquots of each culture were taken and monitored for growth using an Epoch 2 microplate reader (Biotek) in an anaerobic chamber every 15 min until stationary phase was reached. Bacterial growth curves were performed in triplicate with each biological replicate derived from a single isolated colony. Growth curves were plotted using GraphPad.

### RNA sequencing and gene expression analysis

RNA was isolated using TRIzol Reagent (Invitrogen, Cat. No. 15596026) and further processed with the RNeasy Plus Mini Kit (Qiagen, Cat. No. 74034) according to the manufacturers’ protocols. Sequencing reads generated from the Illumina platform were assessed for quality and trimmed for adapter sequences using TrimGalore! v0.4.2 (Babraham Bioinformatics), a wrapper script for FastQC and cutadapt. Reads that passed quality control were then aligned to the mouse reference genome (GRCm38) using the STAR aligner v2.5.1 (Dobin et al., 2013). The alignment for the sequences were guided using the GENCODE annotation for mm10. The aligned reads were analyzed for differential expression using Cufflinks v2.2.1, a RNA-seq analysis package which reports the fragments per kilobase of exon per million fragments mapped (FPKM) for each gene (Trapnell et al., 2010). A differential analysis report was generated using Cuffdiff. Differentially expressed (DE) genes were identified using a significance cutoff of q-value <0.05. The genes were then subjected to gene set enrichment analysis and pathway analysis with iPathwayGuide (AdvaitaBio). Real Time Quantitative PCR Analysis was performed as previously described (Osborn et al., 2021).

### Western Blotting

Whole liver tissue homogenates were made from tissues in a modified RIPA buffer containing protease (Cell Signaling) and phosphatase (Fisher Scientific) inhibitors. Protein was quantified using the bicinchoninic acid assay (Pierce). Proteins were separated by 4–12% SDS-PAGE, transferred to polyvinylidene difluoride (0.45 um, Thermo Scientific) membranes, and proteins were detected after incubation with specific antibodies (see below). Images were captured using a GE Amhersham Imager 6000 and the accompanying software (version 1,1.1) was used for densitometric protein expression analysis. Primary antibodies (CD36, Novus, NB400-144SS; ME-1, Santa Cruz, sc-365891) were prepared 1:1000 in TBST buffer with 5% (w/v) BSA. Secondary (Rabbit HRP, Cell Signaling, #7074; Mouse HRP, Cell signaling, #7076) and β-actin-HRP (Cell Signaling, #12620) antibodies were prepared 1:5000 in TBST with 5% (w/v) non-fat dried milk.

### Liver histology and analysis

Liver histopathology was assessed following preparation based on a previously reported method (Rao et al., 2016) in the Cleveland Clinic Lerner Research Institute Imaging Core. Briefly, following intracardiac perfusion of 10 ml sterile saline via the left ventricle, the liver was removed, weighed, and the right lobe was formalin fixed. After one week of formalin fixation, the right lobe was transferred to a cassette in 70% histology-grade ethanol before embedding in paraffin, sectioning, and staining with hematoxylin and eosin. Liver histopathology was blindly scored by D.S.A. for steatosis, microvesicular steatosis, lobular inflammation, ballooning degeneration, and fibrosis.

### Statistics and reproducibility

Relative abundances were calculated from the raw output of NinjaMap-processed metagenomic data without rarefying the total number of reads across samples. After setting undetected bins to a minimum value of 10^−8^, all relative abundances were further transformed by log10. To evaluate the reproducibility of *in vivo* colonization, Pearson’s correlation coefficients (R) were calculated for the community structure between each two experiments after averaging the relative abundance of each strain across the 3-5 mice that were co-housed in the same cage/experiment. To find the potential strains significantly impacted by strain(s) dropout, multiple unpaired t test with FDR correction was performed for each strain in all mice between two groups. The statistical parameters were set as default with individual variance computed for each comparison followed by multiple comparison of false discovery rate (FDR) with two-stage step-up (Benjamini, K Krieger, and Yekutieli) and Q=0.1%. Dot plots showing relative abundance data were plotted with a conservative lower threshold of 1 × 10^−6^. Further details of statistical analyses can be found in the corresponding figure legends. All statistical analysis and plotting were performed in MATLAB or Prism.

For metabolomic data, metabolite concentration differences between experimental groups or conditions were evaluated using unpaired two-tailed Students’ t test for pairwise comparison, one-way ANOVA for multiple comparisons. All statistical analysis and plotting were performed in Prism.

## ACKNOWLEDGMENTS

We are deeply indebted to members of the Fischbach, Brown, and Hazen labs for helpful discussions. This work was supported by the Human Frontier Science Program LT000493/2018-L (K.N.), a Fellowship from the Astellas Foundation for Research on Metabolic Disorders (K.N.), a research grant from Kanae Fundation for the Promotion of Medical Science (K.N.), the Stanford Microbiome Therapies Initiative (M.A.F.), NIH grants DP1 DK113598 (to M.A.F.), P01 HL147823 (to M.A.F. and J.M.B.), R01 DK101674 (to M.A.F.), P50 AA024333 (J.M.B.), R01 DK130227 (J.M.B.), NSF grant EF-2125383 (M.A.F.) the Bill and Melinda Gates Foundation (to M.A.F.), an HHMI-Simons Faculty Scholars Award (M.A.F.), the Leducq Foundation (M.A.F.), the Leona M. and Harry B. Helmsley Charitable Trust (M.A.F.); and MAC3 Impact Philanthropies (M.A.F.). M.A.F. is a Chan Zuckerberg Biohub Investigator.

## AUTHOR CONTRIBUTIONS

M.W., J.M.B, and M.A.F. conceived and designed the experiments. M.W., A.W., J.Y., L.J.O., W.J.M., V.V., A.H., R.B., D.S.A., A.M.H., A.D., A.Z., K.N., A.G.C., and S.H. performed the experiments. All authors analyzed data. M.W. and M.A.F. wrote the manuscript. All authors discussed the results, commented on the manuscript, and/or added to the manuscript.

