## Supplementary Information for "Strain dropouts reveal interactions that govern the metabolic output of the gut microbiome"

### SUPPLEMENTARY TEXT

#### Technical and biological reproducibility

We constructed hCom1a, a defined 107-member bacterial community (**Figure S1**), by growing its constituent strains separately and then mixing them together without normalization (**Methods**). Cells from the mixed culture were washed, resuspended in 1.2 ml of 25% glycerol, and stored at -80 $^{o}$C.

To test the technical and biological reproducibility of hCom1a colonization, we used three groups of mice colonized by replicates of hCom1a constructed independently on different days. We gavaged germ-free mice with thawed aliquots of the frozen community (**Figure S8A**). After four weeks of colonization, mice were sacrificed and cecal and colonic contents were analyzed by high-resolution metagenomic sequencing (**Table S5**). Strain abundances in all technical replicates were highly similar in the cecum (pairwise Pearson's R=0.97±0.01). The cecal communities in three groups of biological replicates displayed a high degree of similarity in relative abundance profiles (R >0.90 between all pairs, **Figure S8B**). This high degree of reproducibility will enable the use of this complex defined community as an experimental model.

Our analysis of the community composition in all mice as well as the inoculum yielded four conclusions: 1) We confirmed the presence of almost all strains in the inoculum. Of these, 101 strains were detected with a mean relative abundance >1e-6. Two strains showed a low relative abundance (*Anaerofustis stercorihominis* DSM 17244, *Blautia hydrogenotrophica* DSM 10507) and four others were not detected in all three replicates (*Clostridium methylpentosum* DSM 5476, *Dialister invisus* DSM 15470, *Ethanoligenens harbinense* YUAN-3*, Eubacterium dolichum* DSM 3991) (**Figure S8C**). 2) Most strains in the inoculum colonized the mouse gut. 92 strains were detected in the mice cecum at least once; of these, 69 strains were detected with a mean relative abundance >1e-6 (**Figure S8C-D**). 3) Strain relative abundances were tightly distributed in the inoculum but spanned >6 orders of magnitude in the cecum with a coefficient of variation (CV, standard deviation/mean) <0.4 for nearly all strains (**Figure S8C-D**). 4) All 5 bacterial phyla were observed in the cecum. Bacteroidetes dominated, accounting for 83.9% of total reads, followed by Firmicutes (7.7%), Verrucomicrobia (4.1%), Proteobacteria (0.5%) and Actinobacteria (0.05%) (**Figure S8C-D**). Taken together, these data show that hCom1a can colonize GF mice in a reproducible manner, facilitating the strain dropout experiments described in this manuscript.

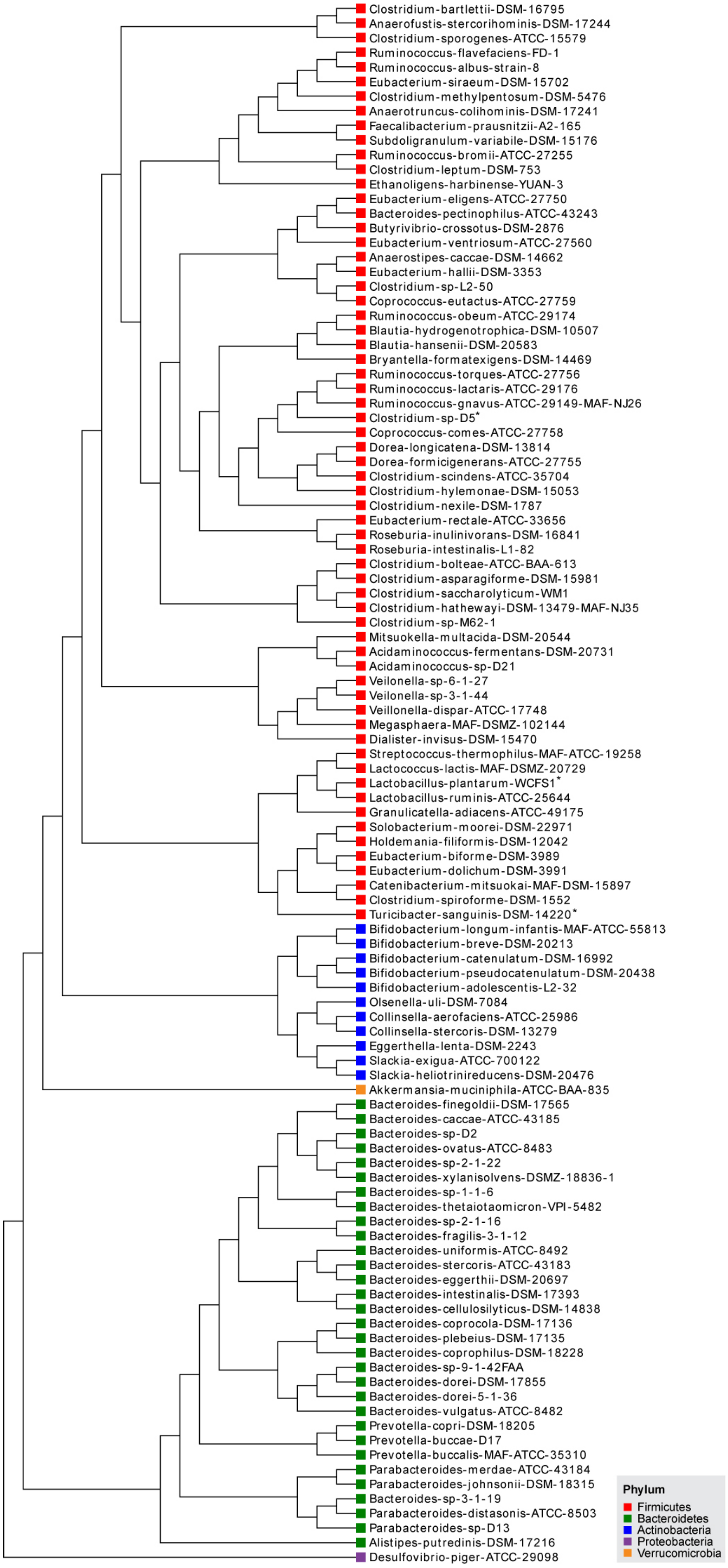
Clostridium-bartlettii-DSM-16795
Anaerofustis-stercorihominis-DSM-17244
Clostridium-sporogenes-ATCC-15579
Ruminococcus-flavefaciens-FD-1
Ruminococcus-albus-strain-8
Eubacterium-siraeum-DSM-15702
Clostridium-methylpentosum-DSM-5476
Anaerotruncus-colihominis-DSM-17241
Faecalibacterium-prausnitzii-A2-165
Subdoligranulum-variabile-DSM-15176
Ruminococcus-bromii-ATCC-27255
Clostridium-leptum-DSM-753
Ethanoligens-harbinense-YUAN-3
Eubacterium-eligens-ATCC-27750
Bacteroides-pectinophilus-ATCC-43243
Butyrivibrio-crossotus-DSM-2876
Eubacterium-ventriosum-ATCC-27560
Anaerostipes-caccae-DSM-14662
Eubacterium-hallii-DSM-3353
Clostridium-sp-L2-50
Coprococcus-eutactus-ATCC-27759
Ruminococcus-obeum-ATCC-29174
Blautia-hydrogenotrophica-DSM-10507
Blautia-hansenii-DSM-20583
Bryantella-formatexigens-DSM-14469
Ruminococcus-torques-ATCC-27756
Ruminococcus-lactaris-ATCC-29176
Ruminococcus-gnavus-ATCC-29149-MAF-NJ26
Clostridium-sp-D5*
Coprococcus-comes-ATCC-27758
Dorea-longicatena-DSM-13814
Dorea-formicigenerans-ATCC-27755
Clostridium-scindens-ATCC-35704
Clostridium-hylemonae-DSM-15053
Clostridium-nexile-DSM-1787
Eubacterium-rectale-ATCC-33656
Roseburia-inulinivorans-DSM-16841
Roseburia-intestinalis-L1-82
Clostridium-bolteae-ATCC-BAA-613
Clostridium-asparagiforme-DSM-15981
Clostridium-saccharolyticum-WM1
Clostridium-hathewayi-DSM-13479-MAF-NJ35
Clostridium-sp-M62-1
Mitsuokella-multacida-DSM-20544
Acidaminococcus-fermentans-DSM-20731
Acidaminococcus-sp-D21
Veilonella-sp-6-1-27
Veilonella-sp-3-1-44
Veillonella-dispar-ATCC-17748
Megasphaera-MAF-DSMZ-102144
Dialister-invisus-DSM-15470
Streptococcus-thermophilus-MAF-ATCC-19258
Lactococcus-lactis-MAF-DSMZ-20729
Lactobacillus-plantarum-WCFS1*
Lactobacillus-ruminis-ATCC-25644
Granulicatella-adiacens-ATCC-49175
Solobacterium-moorei-DSM-22971
Holdemania-filiformis-DSM-12042
Eubacterium-biforme-DSM-3989
Eubacterium-dolichum-DSM-3991
Catenibacterium-mitsuokai-MAF-DSM-15897
Clostridium-spiroforme-DSM-1552
Turicibacter-sanguinis-DSM-14220*
Bifidobacterium-longum-infantis-MAF-ATCC-55813
Bifidobacterium-breve-DSM-20213
Bifidobacterium-catenulatum-DSM-16992
Bifidobacterium-pseudocatenulatum-DSM-20438
Bifidobacterium-adolescentis-L2-32
Olsenella-uli-DSM-7084
Collinsella-aerofaciens-ATCC-25986
Collinsella-stercoris-DSM-13279
Eggerthella-lenta-DSM-2243
Slackia-exigua-ATCC-700122
Slackia-heliotrinireducens-DSM-20476
Akkermansia-muciniphila-ATCC-BAA-835
Bacteroides-finegoldii-DSM-17565
Bacteroides-caccae-ATCC-43185
Bacteroides-sp-D2
Bacteroides-ovatus-ATCC-8483
Bacteroides-sp-2-1-22
Bacteroides-xylanisolvens-DSMZ-18836-1
Bacteroides-sp-1-1-6
Bacteroides-thetaiotaomicron-VPI-5482
Bacteroides-sp-2-1-16
Bacteroides-fragilis-3-1-12
Bacteroides-uniformis-ATCC-8492
Bacteroides-stercoris-ATCC-43183
Bacteroides-eggerthii-DSM-20697
Bacteroides-intestinalis-DSM-17393
Bacteroides-cellulosilyticus-DSM-14838
Bacteroides-coprocola-DSM-17136
Bacteroides-plebeius-DSM-17135
Bacteroides-coprophilus-DSM-18228
Bacteroides-sp-9-1-42FAA
Bacteroides-dorei-DSM-17855
Bacteroides-dorei-5-1-36
Bacteroides-vulgatus-ATCC-8482
Prevotella-copri-DSM-18205
Prevotella-buccae-D17
Prevotella-buccalis-MAF-ATCC-35310
Parabacteroides-merdae-ATCC-43184
Parabacteroides-johnsonii-DSM-18315
Bacteroides-sp-3-1-19
Parabacteroides-distasonis-ATCC-8503
Parabacteroides-sp-D13
Alistipes-putredinis-DSM-17216
Desulfovibrio-piger-ATCC-29098
Phylum
Firmicutes
Bacteroidetes
Actinobacteria
Proteobacteria
Verrucomicrobia

**Figure S1: Phylogenetic tree of hCom1a, the 107-member gut bacterial community used in this study.** The phylogenetic tree was constructed based on a multiple sequence alignment of conserved single-copy genes.

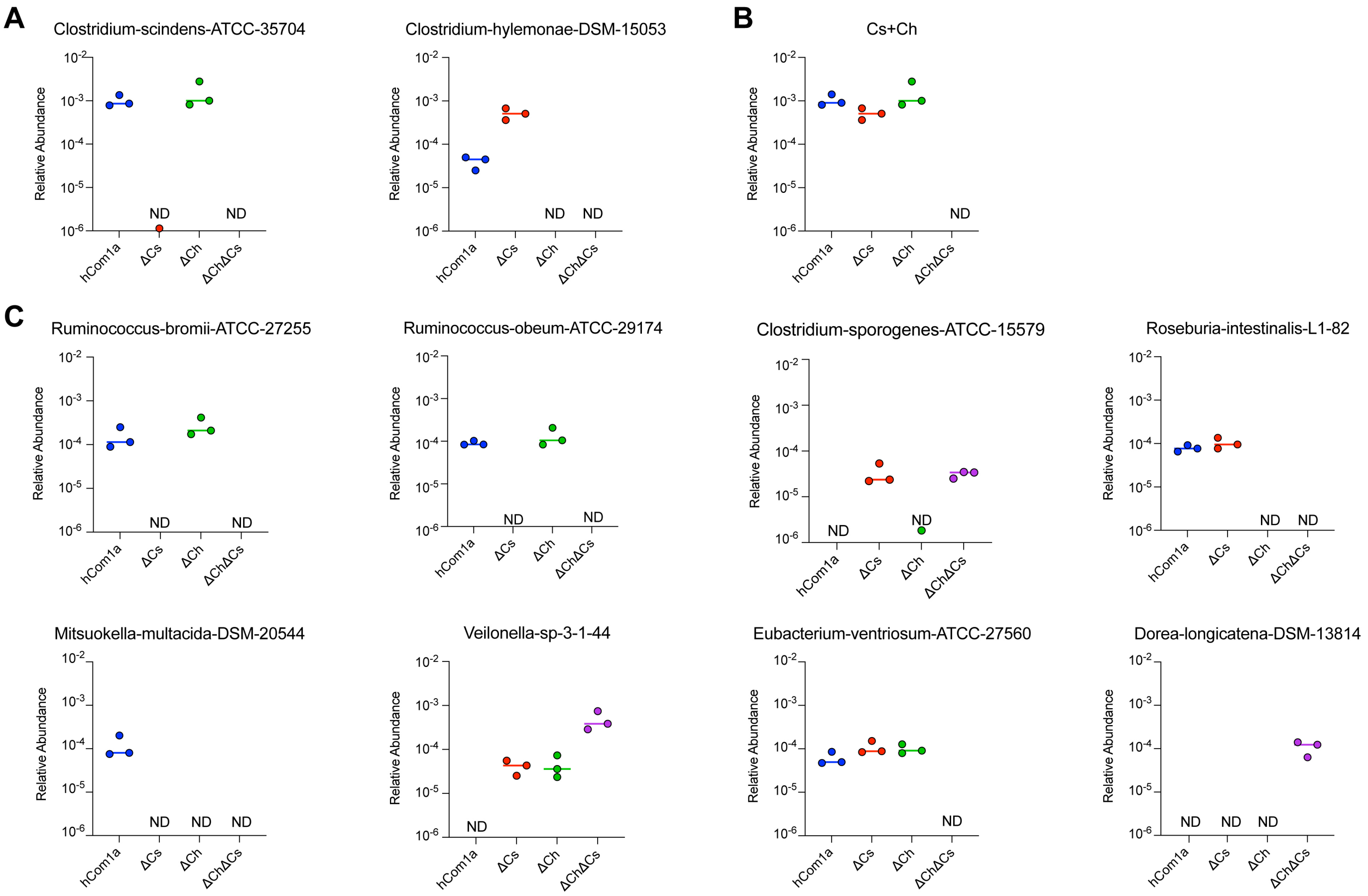


**Figure S2: Relative abundances of select strains in cecal contents of hCom1a-, Δ*Cs*-, Δ*Ch*-, or Δ*Ch*Δ*Cs*-colonized mice.** (**A**) Relative abundances of *C. scindens* and *C. hylemonae.* (**B**) Total relative abundance of *C. scindens* and *C. hylemonae*. (**C**) Relative abundances of the eight interacting strains discovered from the statistical analysis shown in **Figure 2**. n=3 mice per group; ND: not detected (or less than 1e-6).

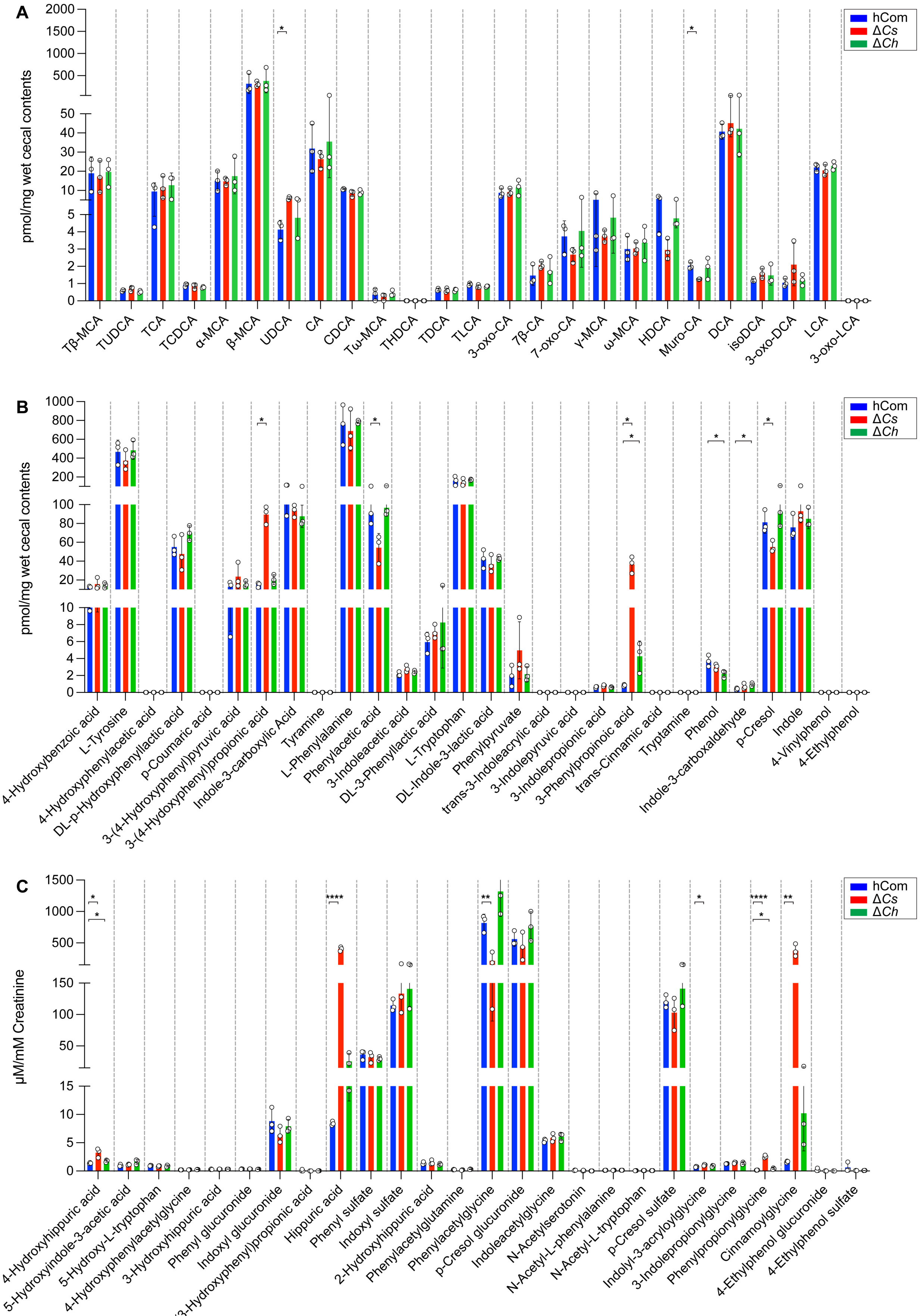
A
pmol/mg wet cecal contents
hCom
ΔCs
ΔCh
Tβ-MCA
TUDCA
TCA
TCDCA
α-MCA
β-MCA
UDCA
CA
CDCA
Tω-MCA
THDCA
TDCA
TLCA
3-oxo-CA
7β-CA
7-oxo-CA
γ-MCA
ω-MCA
HDCA
Muro-CA
DCA
isoDCA
3-oxo-DCA
LCA
3-oxo-LCA
B
pmol/mg wet cecal contents
hCom
ΔCs
ΔCh
4-Hydroxybenzoic acid
L-Tyrosine
4-Hydroxyphenylacetic acid
DL-p-Hydroxyphenyllactic acid
p-Coumaric acid
3-(4-Hydroxyphenyl)pyruvic acid
3-(4-Hydoxyphenyl)propionic acid
Indole-3-carboxylic Acid
Tyramine
L-Phenylalanine
Phenylacetic acid
3-Indoleacetic acid
DL-3-Phenyllactic acid
L-Tryptophan
DL-Indole-3-lactic acid
Phenylpyruvate
trans-3-Indoleacrylic acid
3-Indolepyruvic acid
3-Indolepropionic acid
3-Phenylpropinoic acid
trans-Cinnamic acid
Tryptamine
Phenol
Indole-3-carboxaldehyde
p-Cresol
Indole
4-Vinylphenol
4-Ethylphenol
C
μM/mM Creatinine
hCom
ΔCs
ΔCh
4-Hydroxyhippuric acid
5-Hydroxyindole-3-acetic acid
5-Hydroxy-L-tryptophan
4-Hydroxyphenylacetylglycine
3-Hydroxyhippuric acid
Phenyl glucuronide
Indoxyl glucuronide
3-(3-Hydroxyphenyl)propionic acid
Hippuric acid
Phenyl sulfate
Indoxyl sulfate
2-Hydroxyhippuric acid
Phenylacetylglutamine
Phenylacetylglycine
p-Cresol glucuronide
Indoleacetylglycine
N-Acetylserotonin
N-Acetyl-L-phenylalanine
N-Acetyl-L-tryptophan
p-Cresol sulfate
Indolyl-3-acryloylglycine
3-Indolepropionylglycine
Phenylpropionylglycine
Cinnamoylglycine
4-Ethylphenol glucuronide
4-Ethylphenol sulfate

**Figure S3: Targeted profiling of bile acids (cecum) and aromatic amino acid metabolites (cecum and urine) by LC-MS**. (**A**) Germ-free mice colonized by hCom1a or the single-strain dropout communities (Δ*Cs* and Δ*Ch*) have similar bile acid pools. (**B**) hCom1a- and Δ*Ch*-colonized mice have similar aromatic amino acid metabolites in their cecal contents, while Δ*Cs*-colonized mice are skewed toward reduced aromatic amino acid metabolites (PPA and 4-HO-PPA ↑, PAA and p-cresol ↓). (**C**) hCom1a- and Δ*Ch*-colonized mice have similar aromatic amino acid metabolites in their urine, while Δ*Cs*-colonized mice displayed an altered profile that is skewed toward products of amino acid reduction (hippuric acid and cinnamoylglycine ↑, phenylacetylglycine ↓). Statistical significance was assessed using a Student's two tailed t-test (*: $p<0.05$; **: $p<0.01$; ***: $p<0.001$; ****: $p<0.0001$; n.s.: no significance).

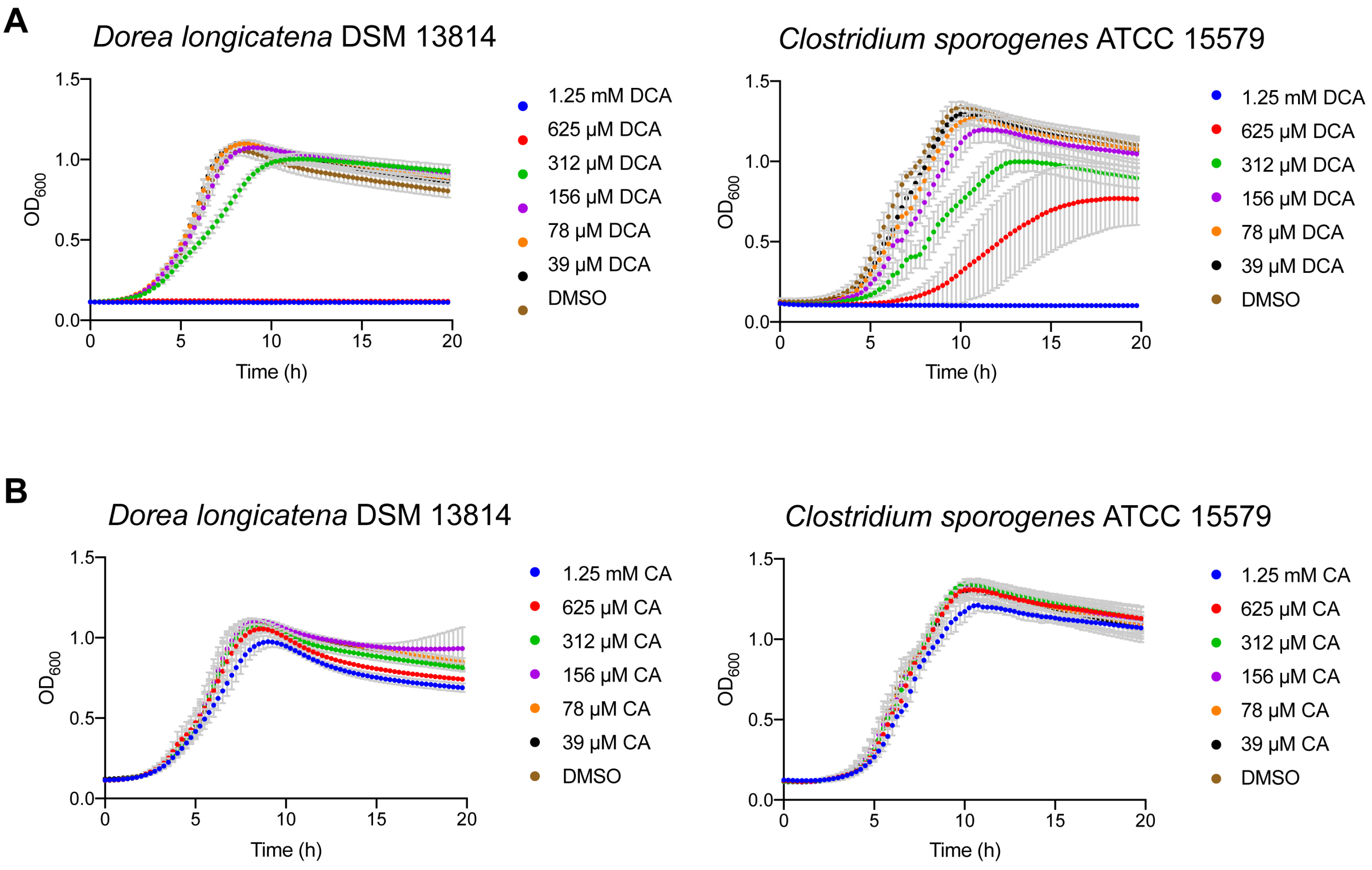


**Figure S4: Growth curves of *Dorea longicatena* and *Clostridium sporogenes* in the presence of various concentrations of cholic acid (CA) or deoxycholic acid (DCA).** (**A**) The growth of *Dorea longicatena* is inhibited by DCA at the physiological concentration of 625 µM, while *Clostridium sporogenes* is only partially inhibited. (**B**) Neither *Dorea longicatena* nor *Clostridium sporogenes* was affected by various concentrations of CA.

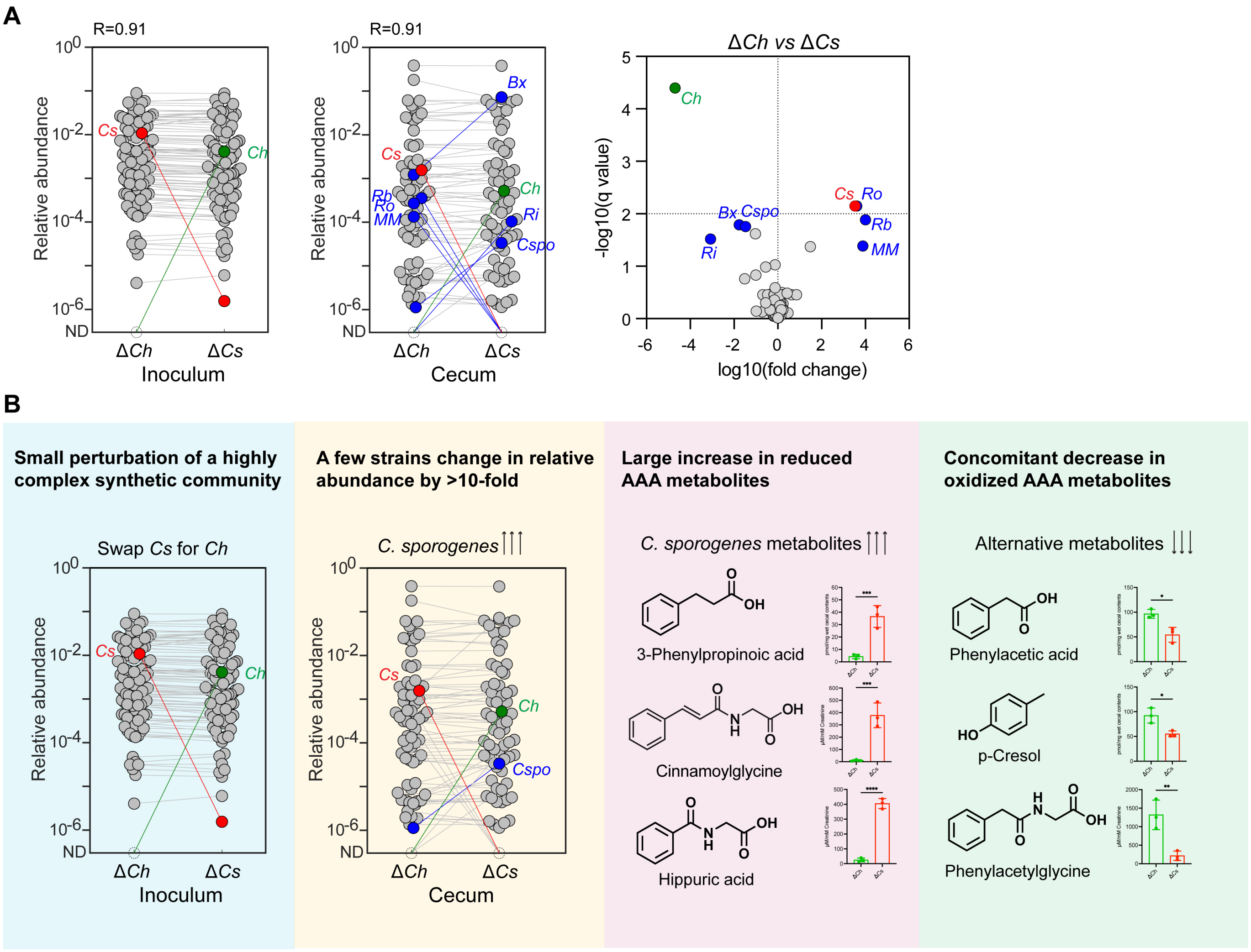


**Figure S5: A strain swap in the 7α-dehydroxylation niche (*Cs* vs *Ch*) has a large impact on community structure and metabolic output.** (**A**) While most strains remain unchanged between the Δ*Cs* and Δ*Ch* dropouts, a small number of strains change in relative abundance >100-fold. *Left:* Metagenomic analysis showing that *Cs* and *Ch* are swapped in the Δ*Ch* and Δ*Cs* community inocula, respectively. *Middle:* The swap between *Cs* and *Ch* impacts the relative abundance of four strains in the Δ*Cs* and Δ*Ch* communities. Each dot is an individual strain; the collection of dots in a column represents the community averaged over 3 mice co-housed in one cage. *Cs* and *Ch* are highlighted in red and green. Strains colored blue went up or down in relative abundance between Δ*Ch* and Δ*Cs*-colonized mice (FDR<0.01, fold change >100). *Right:* Volcano plot showing the log10(relative abundance) values for each strain; for strains that were not detected, relative abundances were set at $10^{-8}$. Strains with significantly different relative abundance (FDR<0.01, fold change >100) are colored blue; the full names of these strains are shown in **Figure 4D**. (**B**) Schematic showing the cascading effects of a single-strain swap within the 7αdehydroxylation niche: the relative abundance of *Cspo* increased from undetectable to ~$10^{-4}$, which increased the abundance of hippurate (~16 fold) and cinnamoylglycine (~37 fold), which decreased the abundance of phenylacetate (~2 fold) and phenylacetylglycine (~6 fold).

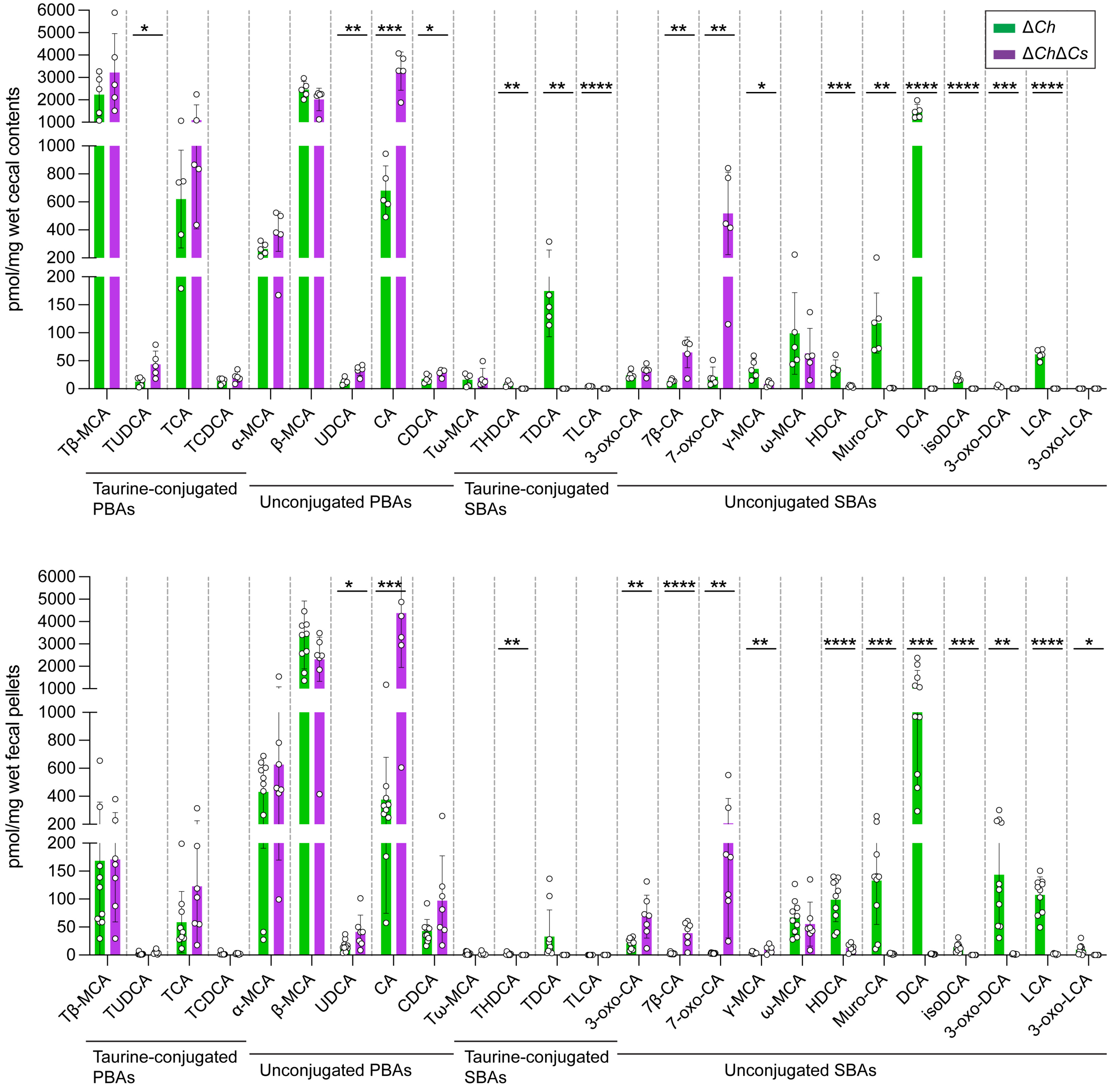


**Figure S6: Targeted bile acid profiling in cecal contents and fecal pellets from Δ*Ch*- and Δ*Ch*Δ*Cs*-colonized mice.** Statistical significance was assessed using a Student's two-tailed t-test (*: p<0.05; **: p<0.01; ***: p<0.001; ****: p<0.0001).

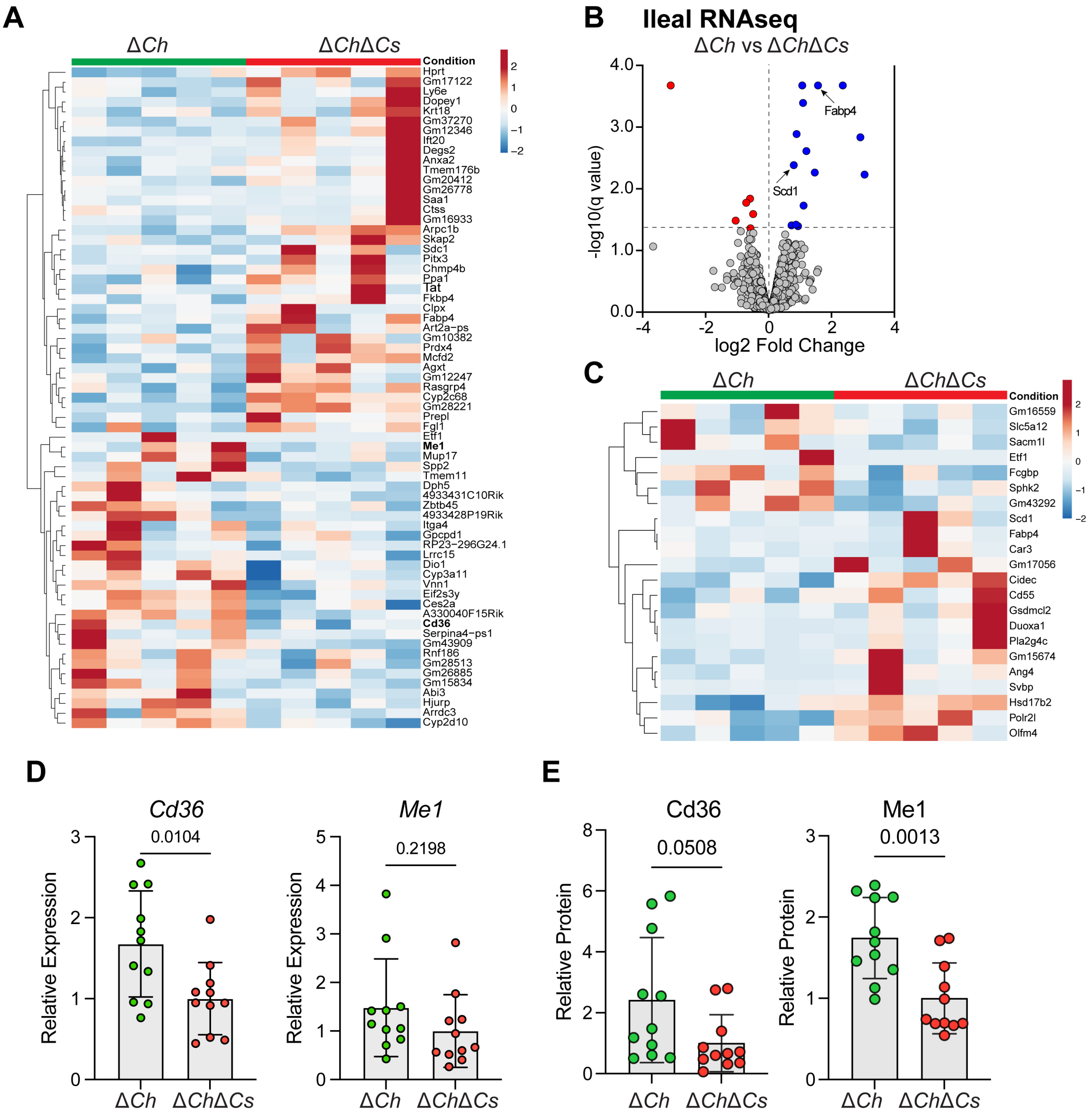


**Figure S7: Dropping out *Cs* has a specific effect on gene expression in the liver and ileum.** (**A**) Heatmap comparing gene expression in liver tissue from Δ*Ch*- vs Δ*Ch*Δ*Cs*-colonized mice. (**B**) Volcano plot of differently expressed genes calculated from RNA-seq of ileal tissue. Genes that are differentially expressed ($q<0.05$) are highlighted in blue (up-regulated) or red (down-regulated). (**C**) Heatmap comparing ileal gene expression in Δ*Ch* vs Δ*Ch*Δ*Cs* groups. (**D**) RT-qPCR results to validate changes in *Cd36* and *Me1* gene expression in the liver. (**E**) Western blot data showing the level of Cd36 and Me1 protein in the liver.

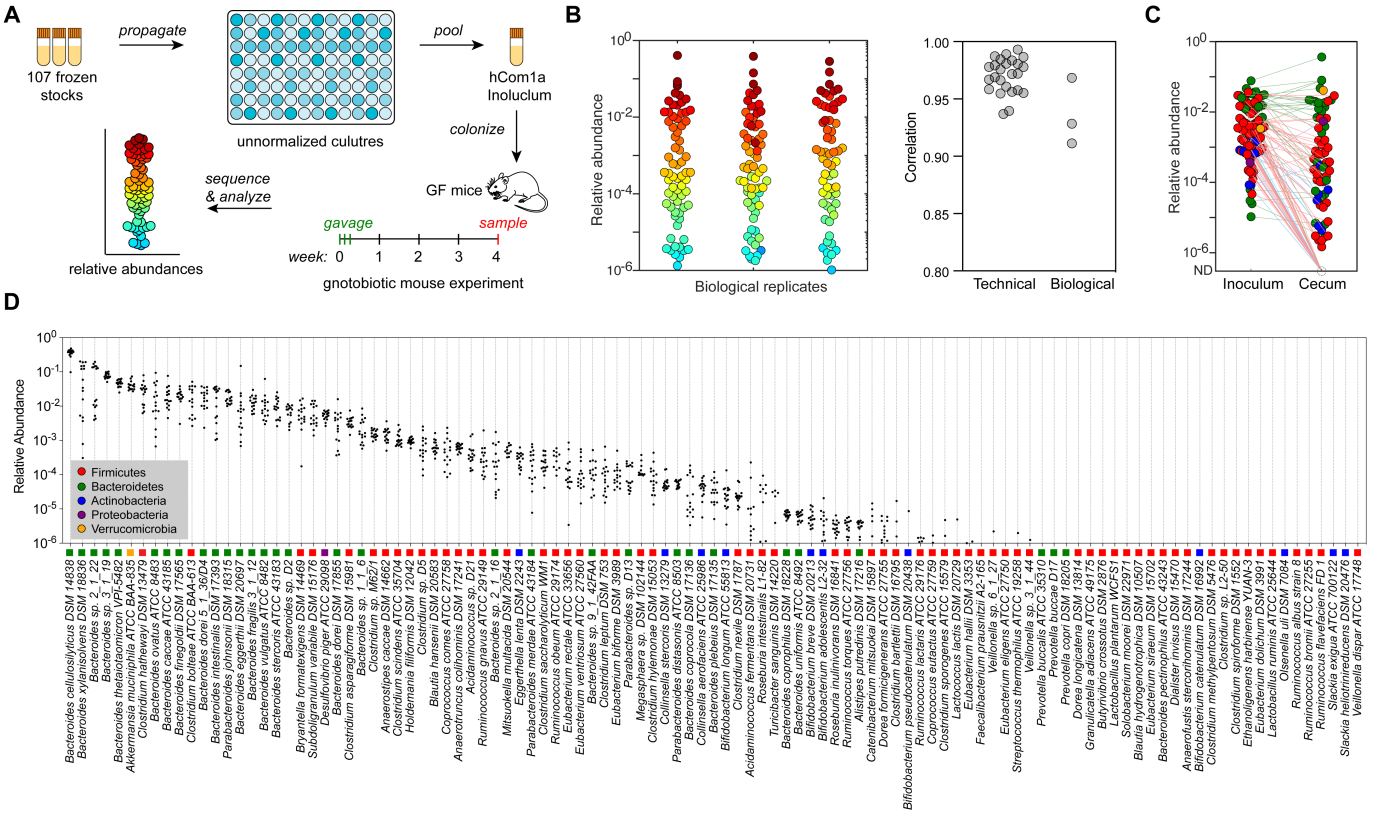


**Figure S8**: **Colonizing germ-free mice with a complex gut bacterial community (hCom1a).** (**A**) Schematic of the experiment. Frozen stocks of the 107 strains were used to inoculate cultures that were sub-cultured every 24 h and then pooled after three days. The mixed culture was used to colonize germ-free Swiss-Webster (SW) mice by oral gavage. After four weeks of colonization, mice were sacrificed and intestinal contents were collected, subjected to metagenomic sequencing, and analyzed by NinjaMap to measure the composition of the community. (**B**) The architecture of hCom1a in the cecum is highly reproducible. Left: community composition is highly similar across three biological replicates. Each dot is an individual strain; the collection of dots in a column represents the community at 4 weeks averaged over multiple mice receiving the same inoculum. Strains are colored according to their average rank-order relative abundance across all samples. Right: Pearson's pairwise correlation coefficients for technical and biological replicates. The log10(relative abundance) values of all strains were used to calculate the Pearson correlation coefficient (R). For strains not detected, the relative abundance was set as 1e-8. (**C**) Averaged relative abundances of the inoculum versus the communities at week 4. Strains in the community span >6 orders of magnitude of relative abundance when colonizing the mouse gut. Dots are colored by phylum according to the legend in panel **D**. (**D**) Relative abundances for most strains are tightly distributed. Each column depicts the relative abundance of an individual strain across all samples at week 4.
